# Chlorine redox chemistry is not rare in biology

**DOI:** 10.1101/2021.12.08.471835

**Authors:** Tyler P. Barnum, John D. Coates

**Affiliations:** Department of Plant and Microbial Biology, University of California, Berkeley, CA 94720, USA

## Abstract

Chlorine is abundant in cells and biomolecules, yet the biology of chlorine oxidation and reduction is poorly understood. Some bacteria encode the enzyme chlorite dismutase (Cld), which detoxifies chlorite (CIO_2_^-^) by converting it to chloride (Cl^-^) and molecular oxygen (O_2_). Cld is highly specific for chlorite and aside from low hydrogen peroxide activity has no known alternative substrate. Here, we reasoned that because chlorite is an intermediate oxidation state of chlorine, Cld can be used as a biomarker for oxidized chlorine species in microorganisms and microbial habitats. Cld was abundant in metagenomes from soils and freshwater to water treatment systems. About 5% of bacterial and archaeal genera contain an organism encoding Cld in its genome, and within some genera Cld is nearly conserved. Cld has been subjected to extensive horizontal gene transfer, suggesting selection by chlorite is episodic yet strong. Cld was also used as a biomarker to predict genes related to chlorine redox chemistry. Genes found to have a genetic association with Cld include known genes for responding to reactive chlorine species and uncharacterized genes for transporters, regulatory elements, and putative oxidoreductases that present targets for future research. Cld was repeatedly co-located in genomes with genes for enzymes that can inadvertently reduce perchlorate (CIO_4_^-^) or chlorate (CIO_3_^-^), confirming that in nature (per)chlorate reduction does not only occur in specialized anaerobic respiratory metabolisms. The presence of Cld in genomes of obligate aerobes without such enzymes suggested that chlorite, like hypochlorous acid (HOCl), might be formed by oxidative processes within natural habitats. In summary, the comparative genomics of Cld has provided an atlas for a deeper understanding of chlorine oxidation and reduction reactions that are an underrecognized feature of biology.

## Introduction

The physical and chemical forms of chlorine are controlled by a biogeochemical cycle ^1^. Chloride (Cl^-^) is the predominant species, and its distribution is largely controlled by physical processes and cellular transport. Organic chlorine species – a diverse range of compounds in which chlorine is a chloro group (-Cl) – are produced and consumed by organisms for chemical defense, signaling, energy, and growth ^1-3^. Inorganic chlorine species – including the chlorine oxyanions hypochlorite (ClO^-^) (and its conjugate acid hypochlorous acid, HOCl), chlorite (CIO_2_^-^), chlorate (CIO_3_^-^), and perchlorate (CIO_4_^-^) – are known to be produced by reduction and oxidation of chlorine ^3-8^. However, a substantial number of questions remain about the extent to these redox reactions participate in biology.

The biology of oxidized chlorine species relates to their high potential to oxidize other molecules. Perchlorate is stable in aqueous solution, but chlorate, chlorite, and hypochlorous acid can be chemically reduced, with each subsequent molecule being more reactive. Reactive chlorine species (RCS) damage cells through oxidative stress ^7,9^. For example, hypochlorous acid causes protein misfolding and sulfur starvation by rapidly oxidizing sulfur in the amino acids methionine and cysteine ^7,9^. Microorganisms from many habitats likely encounter hypochlorous acid ^4,6,7^, atmospherically deposited perchlorate and chlorate ^8^, and other, anthropogenic reactive chlorine species ^10,11^. More biological roles for oxidized chlorine species have been described, including as sources of energy for microorganisms or as chemical weapons ^4,7,12^, but the biology of oxidized chlorines remains incompletely understood. An inventory of habitats in which these chemicals affect organisms, the organisms they affect, and the genes in those organisms potentially involved in chlorine biology would do much to advance our understanding.

The source of oxidized chlorine species within biological habitats depends on the oxidation state of the molecule. Hypochlorous acid can be produced within microbial habitats and cells from chemical or biochemical oxidation of chloride by enzymes like chloroperoxidase ^6,9,13,14 3^. No biological oxidation of chlorine to chlorite, however, has been observed, likely due to the high reduction potential of the redox half-reactions involved (>1 V) ^8^. While (photo)chemical oxidation of aqueous hypochlorous acid to chlorate and perchlorate has been observed experimentally ^15^, in nature production of perchlorate and chlorate is thought to occur predominantly in the atmosphere ^16,17 8^. The diversity of chlorine-oxidizing chemical reactions that occur within biological habitats would be greatly clarified by evidence of which different compounds microorganisms encounter.

The degradation of oxidized chlorine species, aside from hypochlorous acid, is thought to occur predominantly through dissimilatory (per)chlorate reduction, a specialized anaerobic respiratory pathway wherein high affinity perchlorate reductases (Pcr) or chlorate reductases (Clr) reduce perchlorate or chlorate to provide energy in anoxic habitats ^8,18,19^. Reduction may instead occur through co-metabolism: due to the structural and chemical similarity between oxyanions like nitrate and chlorate and perchlorate, enzymes such as nitrate reductase can reduce perchlorate or chlorate ^20-24^. This inadvertent reduction of perchlorate or chlorate produces chlorite and damages cells unless chlorite is degraded ^25^. An unanswered question is if co-metabolic (per)chlorate reduction occurs at a meaningful extent at the low concentrations of perchlorate and chlorate found in nature. If so, many more organisms would contribute to perchlorate and chlorate reduction than presently understood.

Thus, significant gaps remain in our understanding chlorine reduction and oxidation in biology. A promising approach to answer these questions is to identify and use a biomarker for oxidized chlorine molecules. Chlorite dismutase (Cld) is a heme-containing enzyme that catalyzes a chlorite:oxygen lyase reaction wherein a single molecule of chlorite is cleaved into chloride and molecular oxygen, which detoxifies chlorite and yields oxygen ^26-28^. First identified as necessary enzyme in canonical dissimilatory (per)chlorate reducing bacteria ^29,30^, Cld has since been found in bacteria not known to produce chlorite as part of their metabolism ^31^. Subsequent investigations have defined the amino acids required for Cld activity ^32^ and found that aside from low hydrogen peroxidase activity, Cld has no activity towards other compounds, including nitrite, nitric oxide, hydroxylamine, and thiocyanate ^31,33^. These properties make the gene cld a useful, specific biomarker for chlorite. Because chlorite is an intermediate oxidation state of chlorine, organisms encoding Cld in their genomes have likely experienced not only chlorite but also more-oxidized chlorine species that can be reduced to chlorite and more-reduced chlorine oxyanion species to which chlorite is reduced.

Here, we use *cld* as a biomarker for chlorite in microbial genomes to expand what is known about the biology of chlorine oxyanions and redox chemistry. This comparative genomics approach adopts only two assumptions: that organisms encoding Cld experienced chlorite, and that genetic proximity to *cld* means a gene’s product is more likely to function in producing chlorite or responding to its presence. By identifying *cld* and its neighboring genes in thousands of genomes and metagenomes, we were able to describe the distribution of Cld across taxa and environments, expand the evolutionary history of Cld, and predict genes that are functionally related to Cld activity, including biology involving other oxidized chlorine species. These results provide an extensive genomic catalogue for further research in multiple aspects of the biology of chlorine oxidation and reduction.

## Results and Discussion

### Distribution of Cld

Cld proteins belong to the protein family Pfam 06778, which is part of the CDE superfamily ^34^.Non-Cld proteins in Pfam 06778, from which Cld evolved ^35^, are mostly iron-coproporphyrin oxidative decarboxylases (HemQ) that are required for heme biosynthesis in monoderm bacteria ^36^. The use of chlorite dismutase (Cld) as a biomarker requires an accurate definition of proteins with Cld activity, as non-Cld proteins are often incorrectly annotated as Cld or Cld-like proteins in public databases. Here, Cld was defined as proteins in Pfam 06778 that contain the key residues required for Cld activity ^26,32^. Cld proteins formed a monophyletic clade (Figure 1A), confirming previous analyses with smaller datasets ^25,35,37^. Cld proteins were primarily found in diderm phyla (Figure 1A) and were sparsely distributed across the tree of life (Figure 1B). All further investigations of Cld refer only to such proteins, which are further divided into two major clades, lineage 1 and lineage 2 Cld ^38^.

**Figure 1.**
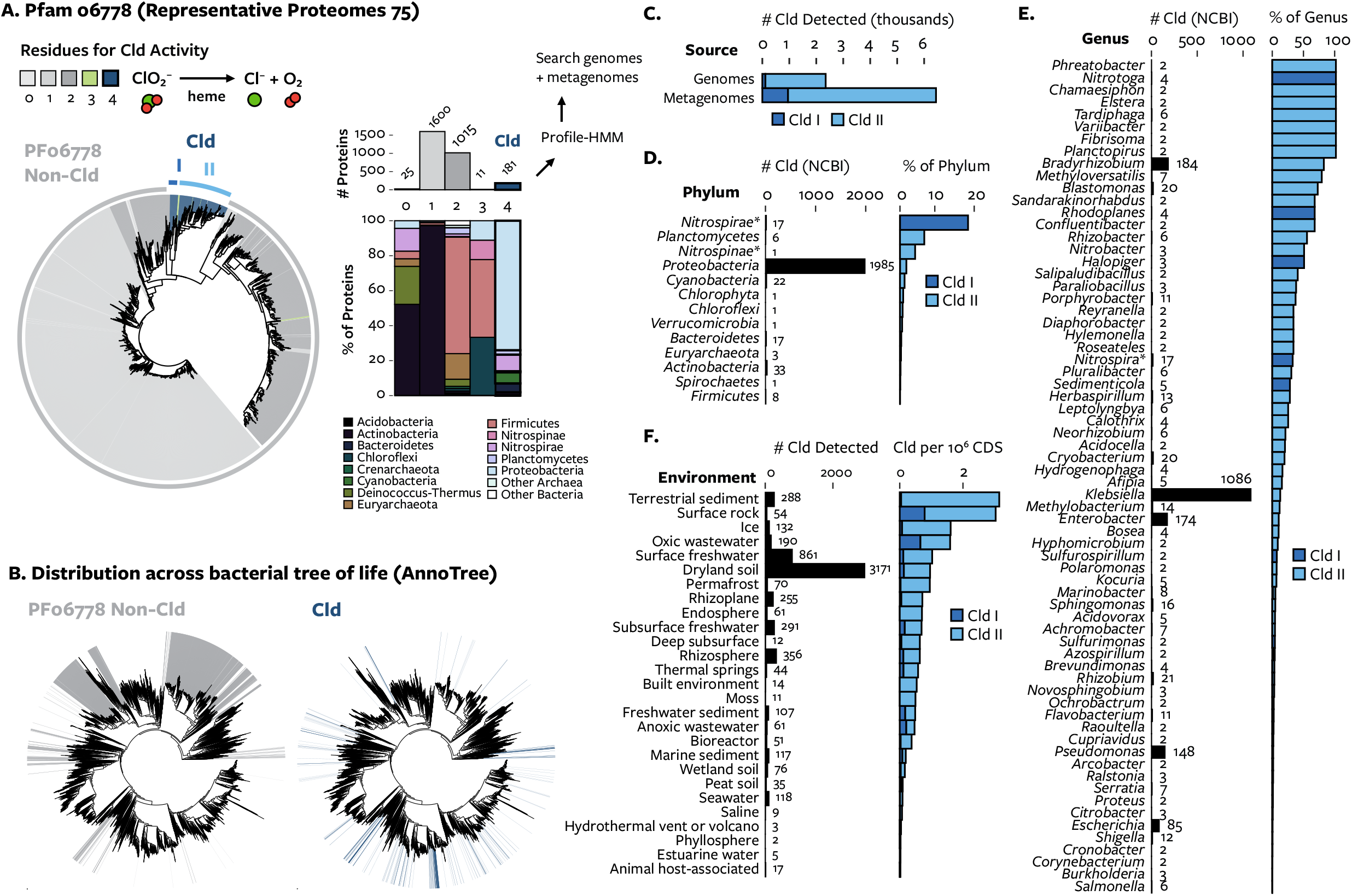
The distribution of Cld across genomes and metagenomes. (A) A maximum-likelihood phylogenetic tree of Pfam 06778, rooted to match Zámocký, et al. ^35^. Color indicates the number of the 4 key residues for Cld activity in each protein. The number of proteins with each fraction of key residues, and the phylogenetic distribution of those proteins, is summarized at right. (B) A tree of all bacterial genomes annotated with the presence Pfam 06778 proteins, comparing the distribution of non-Cld proteins (left, gray) and Cld proteins (right, blue). (C) The total number of each Cld lineage detected in genomes and metagenomes. (D-E) The number (left) and percent (right) genomes within a given RefSeq phylum or genus. For simplicity, only genera with more than one genome encoding Cld and either 20+ genomes or >20% genomes encoding Cld are shown. (F) The number of Cld (left) and fraction of cld per million genes (right) in different environments. Only environments with a sample size of more than 10 million genes are shown. Assuming an average of 5,000 genes per bacterial genome, 1 cld per 1,000,000 genes means that roughly 0.5% of bacterial genomes in a habitat encode Cld.

Profile-HMMs for both lineages of Cld were constructed and used to identify 2411 Cld proteins encoded in 2297 genomes/metagenome-assembled genomes and 6469 Cld in 1575 metagenomes (Figure 1C, Supplementary Data). Here, Cld was identified in 14 phyla and 143 genera, including the bacterial phyla *Actinobacteria, Verrucomicrobia, Firmicutes, Chloroflexi*, and *Spirochaetes* in which Cld has not previously been reported (Supplementary Data). For the first time, Cld was identified in the *Archaea* and *Eukarya*. The low percent identity to bacterial Cld sequences and the similarity of neighboring genes to non-bacterial genes corroborated their assignment to these taxa. The eukaryote with Cld was the unicellular green alga *Monoraphidium neglectum* ^39^. Cld was previously reported in a different eukaryote, the poplar tree (*Populus*) ^38^, but this was later determined to be contamination by bacterial genomic DNA and removed (personal communication, Joint Genome Institute).

Overall, Cld was observed in approximately 1% of genomes, 5% of genera and 15% of phyla in the NCBI taxonomy among the prokaryotes sampled. Genomes from the phyla *Nitrospirae, Planctomycetes*, and *Nitrospinae* are most likely to contain Cld, followed by *Proteobacteria* and *Cyanobacteria* (Figure 1D). The frequency of Cld in *Nitrospirae* and *Nitrospinae* may be underestimated due to the large number of incomplete metagenome-derived genomes in these phyla. At the genus level, typically only a fraction of genomes had Cld, although Cld could be highly conserved within a genus (Figure 1E). Many of these genera belong to specific biological groups, like symbiotic nitrogen-fixing bacteria or nitrite-oxidizing bacteria, but it is unknown whether Cld is related to these biological functions. The extent to which Cld can be found outside of dissimilatory perchlorate- and chlorate-reducing bacteria, and potentially associated with other specific microbial lifestyles, far exceeds that described previously.

The widespread nature of Cld was further supported by its distribution across a dataset of 6961 IMG/M metagenomes encoding 10.8 billion genes. *cld* were a very low proportion of genes in host-associated systems and a greater proportion in freshwater and soil systems (Figure 1F). Comparing metagenomes with greater than 10 million genes with this metric indicated that *cld* was most enriched in environments such as oligotrophic rocks, sediment, and ice followed by oxic wastewater, surface freshwater, and dryland soils. Curiously, within aquatic environments, *cld* appears least frequently where chlorine is most concentrated: estuary, ocean, and hypersaline waters. The shared features of environments where *cld* is the highest proportion of coding genes is that they are predominantly oxic, and many are exposed to high amounts of sunlight ^40,41^. The broad distribution of the *cld* gene in genomes and metagenomes shows that chlorite is experienced by many organisms in a variety of environments. Instead of being a tool of specialized anaerobic metabolisms found in anoxic habitats, Cld is encoded by various organisms from both oxic and anoxic habitats.

### Evolution of Cld

The evolutionary history of Cld may help clarify its biological role. The phylogeny of Cld could be more thoroughly defined using this expanded set of genomic and metagenomic proteins. A phylogenetic tree of 8924 Cld was constructed, grouped into 60 clades with a median clade size of 11 proteins and a maximum clade size of 2936 proteins, and annotated with data about the Cld protein (Figure 2). Only about half of the sequence diversity of Cld is present in cultured organisms (Figure 2A). There is no indication in these data that Cld proteins have lost chlorite:O_2_ lyase activity: many of the largest clades include Cld proteins with biochemically verified activity, and the key residues for chlorite:O_2_ lyase activity were conserved in all clades except the under-sampled clades 10 (n=2) and 32 (n=1) and clade 38 (n=21) (Figure 2B).

**Figure 2.**
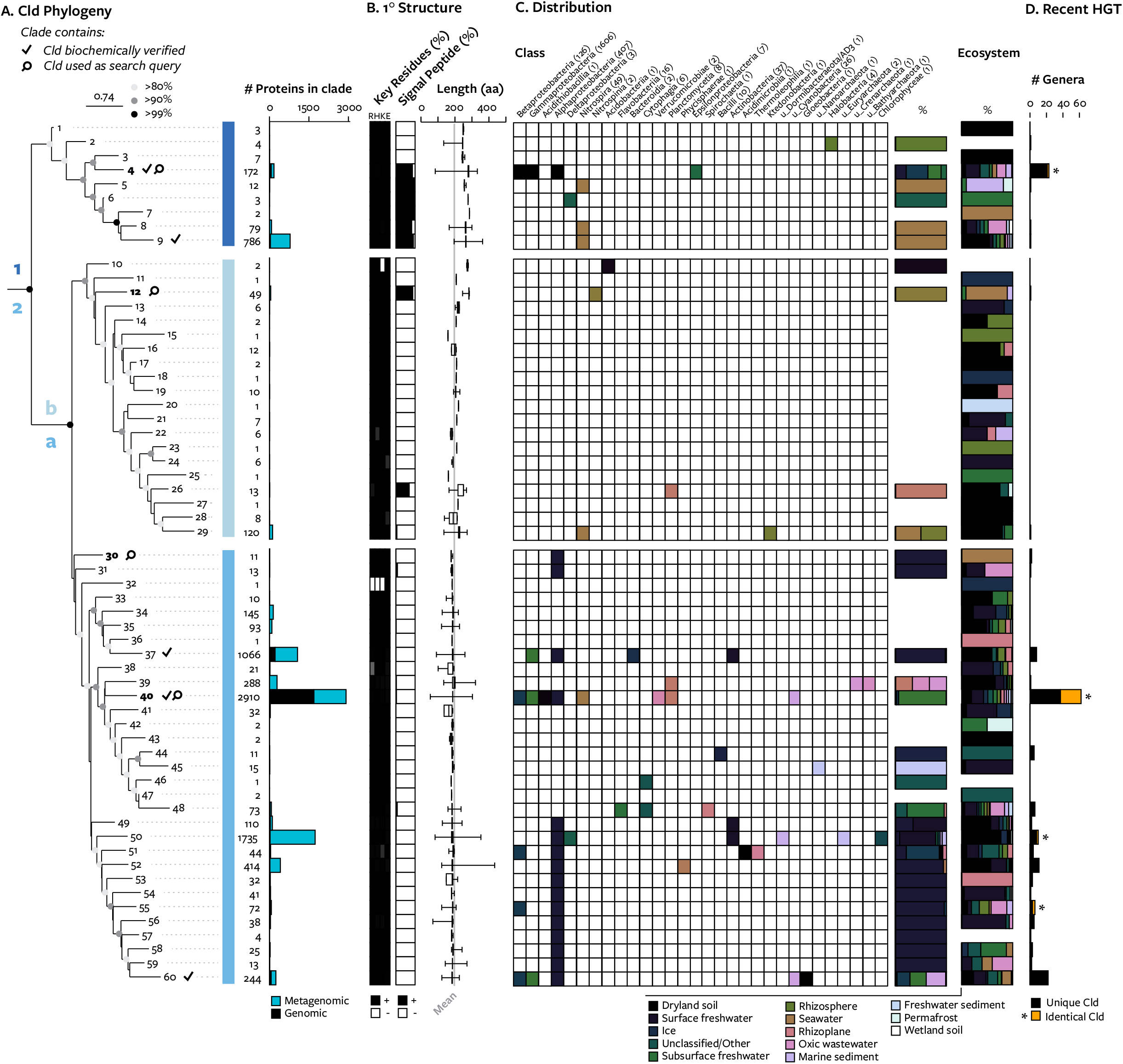
The phylogeny of Cld proteins and attributes of each lineage. (A) A maximum likelihood phylogenetic tree of Cld, with clades formed by phylogenetic distance and node support values indicated by color. Clades containing biochemically verified Cld proteins are indicated by a checkmark, and clades containing sequences used in the initial search are indicated by a magnifying glass. Major lineages of Cld are demarcated by dashed lines. The number of Cld per clade is listed at right and represented in a barplot by whether the source is genomic and metagenomic. (B) The primary structure of each clade. The proportion of each Cld clade computationally identified to have each key residue from 0% (white) to 100% (black) (left); the proportion of the clade computationally predicted to have a signal peptide (black) for export to the periplasm (center); and length of proteins in the clade represented as a boxplot where the box represents the interquartile range, whiskers represent maximum and minimum values, and the gray line represents the mean of all Cld (right). (C) Detection of Cld in each taxonomic class indicated by filled squares (left). Stacked barplots represent the proportion of each Cld clade in each environment (left) or each taxonomic class (right). (D) A measure of recent horizontal gene transfer: the number of genera found in each clade, with genera that can share an identical copy of Cld colored orange.

These data do revise the previous understanding of Cld being composed of two lineages wherein lineage 1 Cld are larger, periplasmic proteins and lineage 2 Cld are smaller, cytoplasmic proteins ^26,42^. First, tree topology showed two distinct, diverse, and strongly supported (>99% bootstraps) sublineages within lineage 2 Cld, which we term lineage 2a and 2b. The only cultivated organism with lineage 2b Cld is *Nitrospina gracilis* ^43^. lineage 2b proteins are an intermediate length of 229 amino acids, and considerable variation in protein size was observed within the shorter lineage 2a Cld (Figure 2C): 4% of Cld had larger (>20 aa) N- and C-terminal extensions that could either be artifacts of protein prediction or fusion proteins that augment the function of Cld ^44^. Second, signal peptide prediction suggested that the more basal branching clades of group 1 Cld are not periplasmic, while two clades of lineage 2b Cld are periplasmic and a small number of Cld from various lineage 2a clades are periplasmic (Figure 2C). This indicates a general purpose of Cld for degradation of intracellular chlorite but periodic selection for the degradation of extracellular chlorite through the acquisition of peptide signals for export, in any lineage.

Horizontal gene transfer of Cld across evolutionary time is evident from its taxonomic distribution. A single taxonomic group can have organisms with Cld from different clades (Figure 2C). A single Cld clade can be found in taxonomic groups spanning phyla or even domains of life. Cld has even been subject to recent transfer between genera: multiple Cld clades consisted of Cld from different genera (Figure 2D). Alone, this metric reflects the combined signal of vertical and horizontal inheritance, but a detailed view shows that horizontal inheritance is a large component. For example, within clade 4, there are two instances where Cld from (per)chlorate-reducing proteobacteria appeared to have been acquired by nitrite-oxidizing *Nitrotoga* (Supplemental Figure 2) ^45^. Representing the most recent horizontal gene transfer: genomes from different genera possessed identical Cld proteins (Figure 2D). In one case the same Cld protein (WP_011514928.1) was found in genomes from 18 genera and spanning *Alphaproteobacteria, Gammaproteobacteria, Betaproteobacteria*. One remarkable recent horizontal gene transfer is the acquisition of periplasmic Cld by *Nitrosomonas mobilis* Ms1, which was isolated from a wastewater treatment plant that used chlorine-based disinfectants (personal communication, Hirotsugu Fujitani) ^46,47^. That Cld has never been observed in ammonia-oxidizing microorganisms prior to this is evidence of how the evolution of Cld is now also being shaped by anthropogenic sources of chlorite.

The scope of horizontal gene transfer and the degree to which it has occurred suggests occasional yet strong selection for the ability to degrade chlorite. The apparent benefit of Cld and ease of its horizontal gene transfer can be reconciled by its rare conservation within phylogenetic groups (Figure 1) by invoking a selective pressure for loss of Cld, possibly related to the heme requirement for this enzyme.

### Comparative Genomics of Cld

Diverse organisms have experienced enough chlorite to select for the cld gene (Figures 1-2). Because genes located near each other in and across genomes are more likely to be functionally related ^48,49^, co-location of genes with *cld* across genomes provides an opportunity to identify other genes likely to be involved in biological processes involving chlorite. We refer the set of genes within 10 genes upstream and downstream of cld as a “genomic neighborhood.” 8,751 genomic neighborhoods contained 61,215 proteins that were clustered into 11,081 protein subfamilies. Subfamilies with a low “clustering coefficient” (Methods, Figure 3A) are found in many different types of genomic neighborhoods with Cld and, therefore, are more likely to have a function related to chlorine redox biology, rather than be co-located by chance.

**Figure 3.**
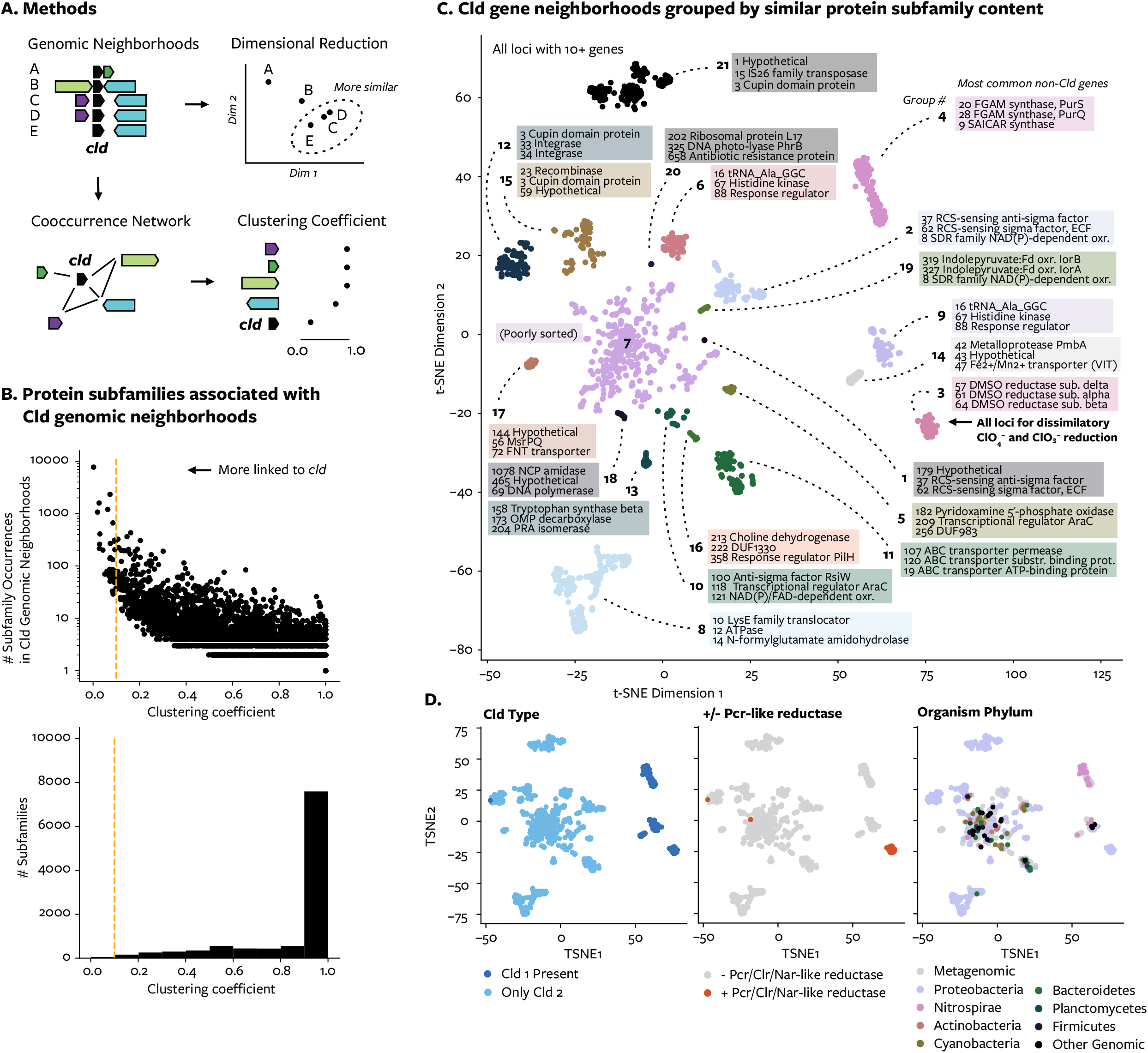
Statistical analysis of Cld genomic neighborhoods. (A) Schematic diagram explaining analyses. Genomic neighborhoods are compared using proteins clustered into protein subfamilies. In the gene-centric analysis, the co-occurrence of genes in different neighborhoods is used to construct a network, from which a clustering coefficient for each gene is derived. In the neighborhood-centric analysis, neighborhoods with more similar gene content are plotted through several dimensional reduction steps and clustered. (B) The distribution of protein subfamilies by their clustering coefficient, a measure of linkage to *cld*. The threshold value for defining “hits” is indicated. (C) Cld genomic neighborhoods colored by group, indicating thee top three most common proteins subfamilies in each group. Genomic neighborhoods that did not cluster into distinct groups are found in neighborhood group 7. (D). Cld genomic neighborhoods colored by the presence of group 1 Cld (left), by the presence or absence of reductases closely related to Pcr, Clr, and Nar (center), or by the phylum of the host organism (right).

Only a small fraction of protein subfamilies in Cld genomic neighborhoods showed a genetic correlation with Cld (Figure 3B). Of most interest are proteins with the lowest clustering coefficients (Table 1), which have the strongest genetic association to Cld. Among these proteins were proteins with functions already known to be connected to Cld, such as (per)chlorate reductases, oxidative stress response, genetic mobility, and signaling ^50-52^. Additionally, since this work began, an alkylhydroperoxidase AhpD-like protein (subfamily 84), identified here as having a low clustering coefficient (Table 1), was found to be the enzyme RcsA involved in hypochlorous acid degradation in Pseudomonas aeruginosa ^53^. Therefore, this method identified true genetic associations between Cld and other protein subfamilies.

**Table 1.**
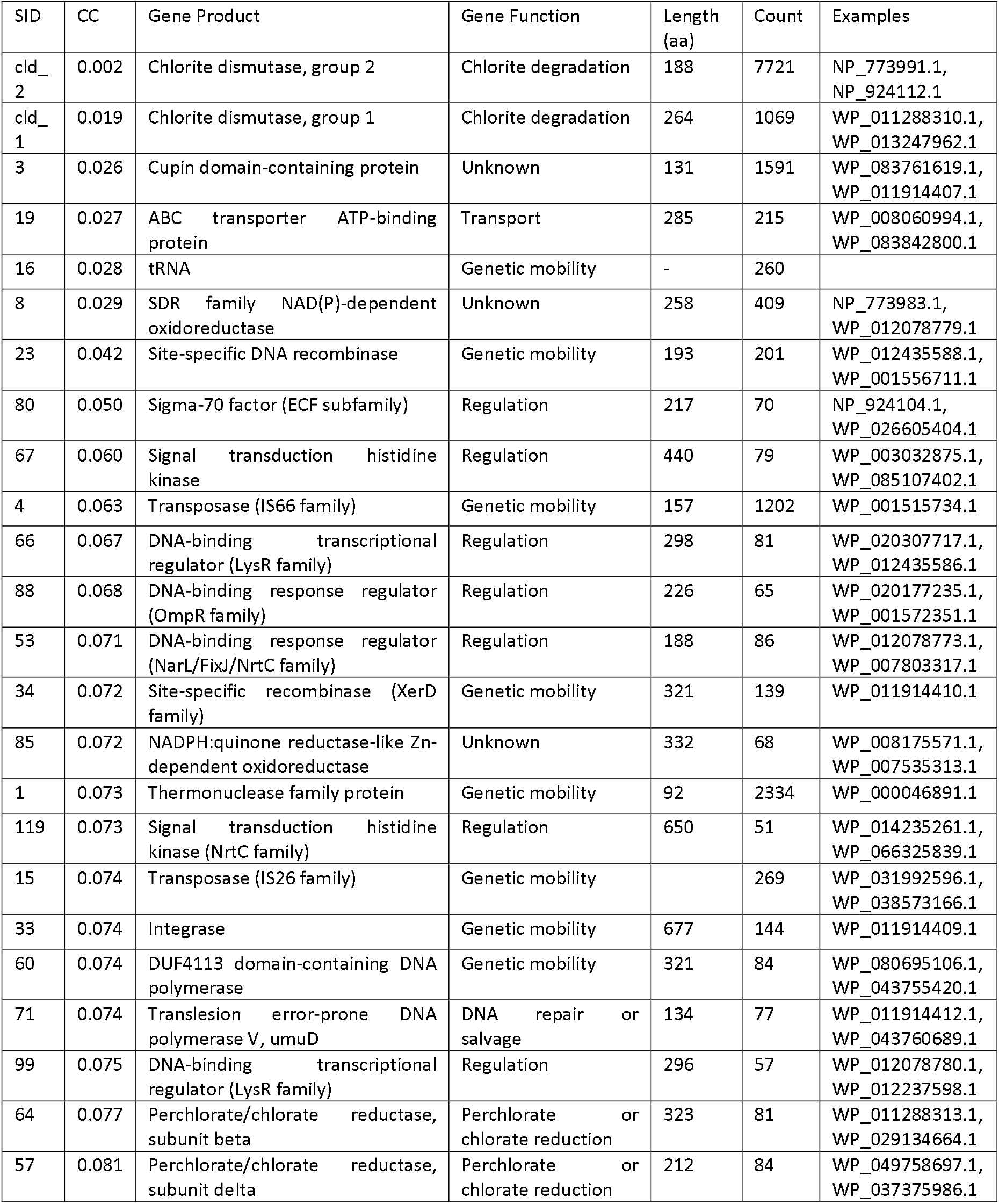
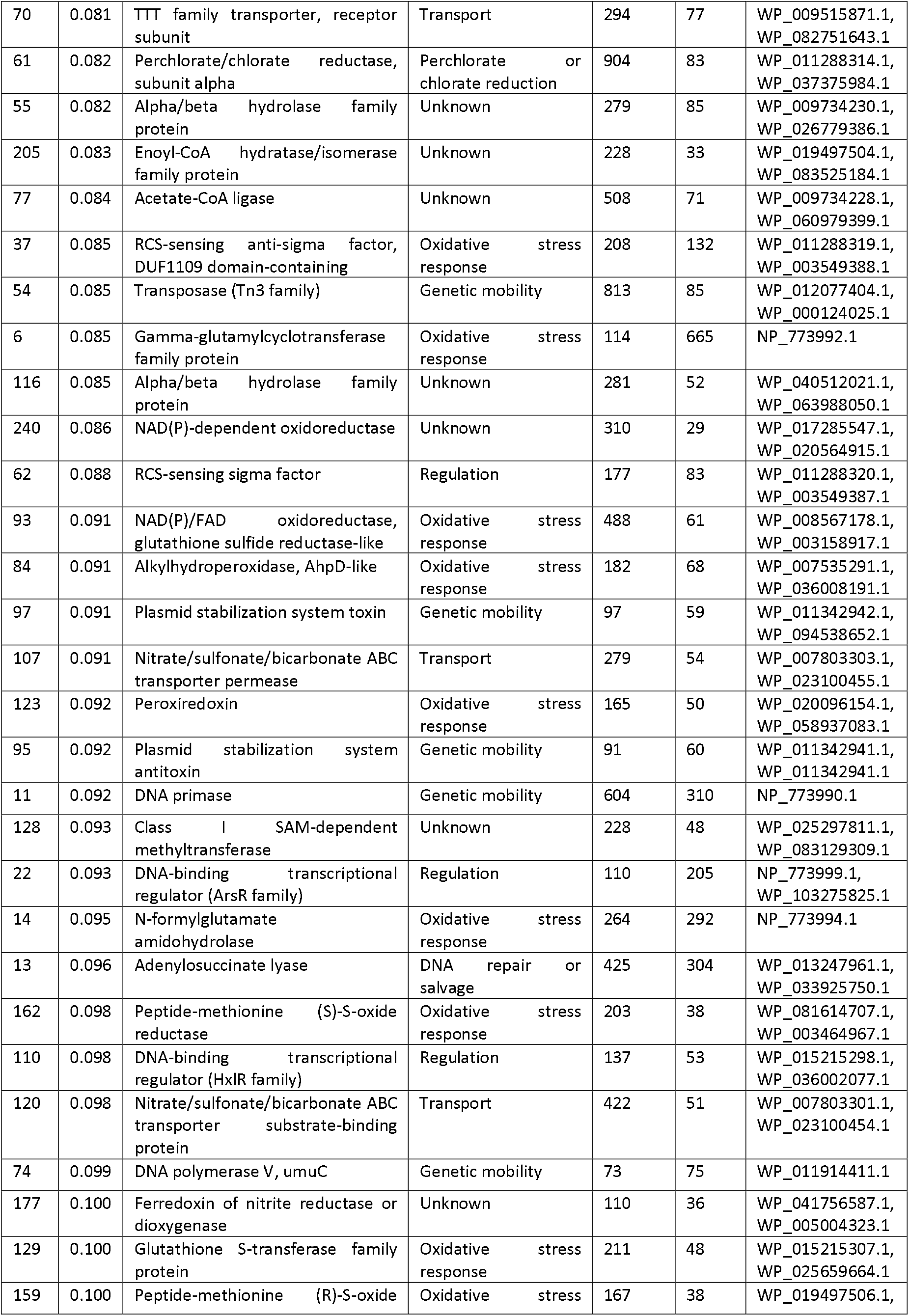
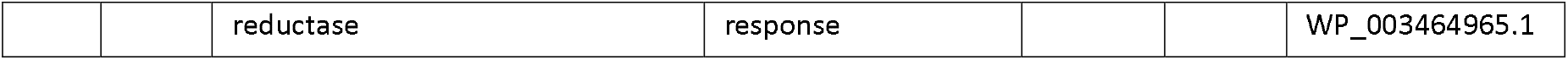
Genetic linkage of protein subfamilies to Cld. All subfamilies with a network clustering coefficient of less than 0.1 are shown. Columns: SID, subfamily ID; CC, clustering coefficient; Gene Function, the predicted role for the gene in chlorine oxyanion biology; Length (aa), mean protein length; Count, total number of genes in the in the subfamily found in Cld genomic neighborhoods; and Examples, RefSeq accession for proteins in the subfamily from different classes of organisms.

Groups of highly similar genomic neighborhoods were defined by unsupervised clustering of neighborhoods with greater than 10 genes (Figures 3A). Neighborhood groups were distinguished by their most abundant non-Cld protein subfamilies (Figure 3C) and recapitulated major differences in Cld (e.g. group 1 vs. 2 Cld, presence or absence of Pcr/Nar-like reductases) (Figure 3C). In total, 20 distinct genomic contexts were produced by clustering, and more groups would likely result with the inclusion of more genomes and metagenomes. Emblematic of this functional diversity is that all known genomic islands and composite transposons for respiratory perchlorate and chlorate reduction – the only biological pathway Cld has been confirmed participating in naturally – were contained in one single neighborhood group (Clark et al 2013, Melnyk and Coates 2015).

Clustering coefficients for protein subfamilies and grouping of neighborhoods allows exploration of relationships between protein families. Among the most interesting genetic associations to Cld were those of putative oxidoreductases with unknown function. Such oxidoreductases accounted for many of the subfamilies with the lowest clustering coefficients (Table 1): cupin domain-containing protein (subfamily 3), NADPH:quinone reductase-like Zn-dependent oxidoreductase (subfamily 85), and SDR family NAD(P)-dependent oxidoreductase (subfamily 8). Neighborhood-group 2 contained the SDR family NAD(P)-dependent oxidoreductase ^54^ as well as reactive chlorine-sensing regulatory elements that also had low clustering coefficients (subfamilies 37 and 62), further implicating a role this subfamily in reactive chlorine stress response. Curiously, fitness data for a protein in *Sphingomonas koreensis* DSMZ 15582 with 47% amino acid identity to a related SDR family NAD(P)-dependent oxidoreductase (subfamily 827, clustering coefficient 0.15) showed a deleterious effect when this protein was disrupted only in chlorite stress conditions or when glutamic acid was the carbon source ^55^. The cupin domain protein was one of the most common subfamilies in the dataset, being found with Cld in 1487 genomes among 40 genera. In fact, encoded with transposases in neighborhood groups 12, 15, and 21, the cupin domain protein could be found in 90% of genomic neighborhoods with the most extreme form of horizontal gene transfer: encoding Cld proteins that are identical across different genera. While the cupin domain protein has been suspected to have a role in reactive chlorine species response in (per)chlorate reducing bacteria ^56^, these data point to a far more common and important role in chlorine redox biology.

In addition to providing genes and loci for reverse genetics, these genetic associations are useful for further describing chlorine redox biology. Cld genomic neighborhoods were first searched for genes described as participating in the response to reactive chlorine species: methionine sulfoxide reductases, sulfur homeostasis proteins, protein chaperones, regulatory systems, and scavenging of reactive byproducts like peroxides, aldehydes, and glyoxals ^7,56,57^. One or more of these genes could indeed be found in Cld neighborhood-groups (Supplemental Data), and Cld was routinely found with methionine sulfoxide reductase systems (Supplemental Figure 3). This confirms that organisms use these genes are not experimental artifacts but respond to reactive chlorine species in nature. Additionally, it provides more evidence that chlorite is produced at a sufficient flux in the environment to contribute to oxidative damage in microorganisms.

The transport of chlorine oxyanions across the cellular membrane appeared to be a defining feature of two types of Cld genomic neighborhoods (Figure 3C). No specific transporters for chlorine oxyanions are known. Neighborhood group 11 contained by ABC transporter subunits, some of which are annotated as ATP-driven nitrate transporters. Such transporters could be involved in the transport of chlorate, a structural analogue of nitrate, an activity previously identified for nitrate transporters by genetic selection for chlorate resistance for example, see: ^58^. Neighborhood group 17 was distinguished by a formate-nitrite transporter (FNT) family protein, an MsrP protein involved periplasmic reactive chlorine stress response (see below), and cytoplasmic Cld. As with nitrate and chlorate, formate (HCO_2_^-^) and nitrite (NO_2_^-^) are structural analogues of chlorite (ClO_2_^-^), and the potential for FNT family proteins to transport chlorite as well has been shown by the deleterious nature of FocA formate transporters and NirC nitrite transporters in chlorite stress conditions ^55^. Curiously, the FNT-Cld-MsrP gene cluster belonged to metagenomic *Mycobacteria* found in seasonally low-oxygen lakes ^59,60^. The combination of a chlorite-permeable transporter and cytoplasmic Cld might act to import extracellular chlorite to be converted to oxygen inside the cell. Microorganisms benefitting from the production of oxygen by Cld is a trait thus far observed only in (per)chlorate-reducing bacteria or engineered strains ^8,61^.

Chlorination and dechlorination are only known to be related to hypochlorous acid, not higher oxidation states of chlorine like chlorite. Relatively low clustering coefficients with Cld for two protein subfamilies suggested otherwise: non-heme chloroperoxidase (subfamily 122), which chlorinates organic molecules by producing hypochlorous acid, and a putative subfamily of haloacid dehalogenases (subfamily 172), which removes chlorine from organic molecules. The relationship could be that chlorite produces and may be produced by hypochlorous acid, which generates stable chlorinated products like chlorotyrosine ^62^, and microorganisms use dehalogenases to reverse that chlorination.

### Chlorite from Chlorine Reduction

The involvement of the above biological functions in chlorine redox biology may be closely related to how chlorite is produced. The major known source of chlorite in biology is the enzymatic reduction of perchlorate and chlorate. To determine which biochemical pathways contribute to chlorine reduction, Cld genomic neighborhoods were searched for proteins in the DMSO reductase family of molybdopterin enzymes (Pfam 00384), which includes the respiratory perchlorate and chlorate reductases (Pcr, Clr) and the enzymes that might inadvertently/co-metabolically reduce those molecules while acting in other biochemical pathways ^20^. If the enzyme reduces perchlorate or chlorate to chlorite in nature, Cld can provide a benefit by degrading chlorite, and the selective pressure to co-express the reductase with would lead to their genetic co-location in some genomes (Figure 4A).

**Figure 4.**
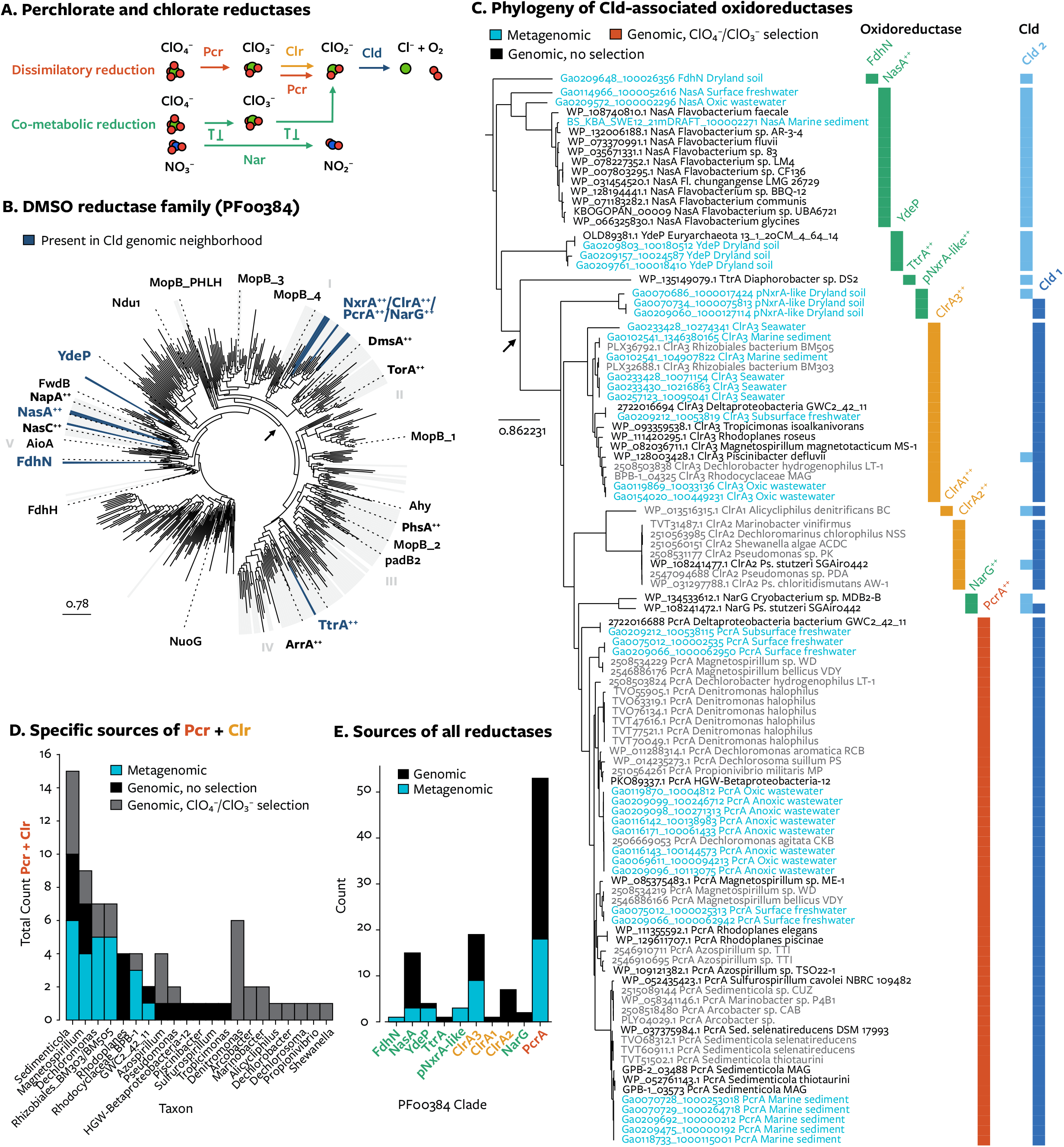
The distribution of Cld among possible perchlorate and chlorate reductases in the DMSO reductase family. (A) The pathways by which a reductase can produce chlorite, which Cld degrades. Dissimilatory reduction occurs through perchlorate reductase (red, Pcr) or chlorate reductase (orange, Clr). Co-metabolic reduction (green) does not occur through a reductase specialized for perchlorate or chlorate reduction. An example is shown for nitrate reductase (Nar). (B) An unrooted maximum likelihood phylogenetic tree of representative proteins from the DMSO reductase family. Clades containing proteins from Cld genomic neighborhoods are highlighted in blue. (C) The same phylogenetic tree omitting all proteins not found with Cld. Colors indicate their source (genomic or metagenomic). Labels at right indicate the type of protein and the lineages of Cld present in their genomic neighborhood. (D) The number of genomes per genus or other taxon with proteins from PcrA, ClrA1, ClrA2, or ClrA3, and whether or not the organisms were subjected to selection for those genes (i.e. providing perchlorate or chlorate as a sole respiratory electron acceptor). Metagenomic proteins were assigned to the closest genomic relative’s taxon. (E) The number of Cld-associated proteins in each clade of the DMSO reductase tree and whether they were obtained from genomes and metagenome-assembled genomes (black) or metagenomes (blue).

A total of 105 proteins in the DMSO reductase family were found in Cld genomic neighborhoods. Many genomic neighborhoods contained cytoplasmic Cld and enzymes with documented in vitro (per)chlorate reductase activity (Figure 4B-E): assimilatory nitrate reductases (NasA) of three phylotypes, a cytoplasmic dissimilatory nitrate reductase (NarG) from a *Cryobacterium* genome ^63^, and a tetrathionate reductase (TtrA) from a *Diaphorobacter* genome ^64^. The co-occurrence of Cld with formate dehydrogenase (FdhN) and an uncharacterized Fdh-like protein (YdeP) was unexpected but could be related somehow to the structural similarity of chlorite and formate. Cld was also found on soil metagenome contigs with uncharacterized enzymes most similar to periplasmic nitrite oxidoreductases (pNxr) of nitrite-oxidizing bacteria (*Nitrospira* and *Nitrotoga*), anammox bacteria, and other organisms ^45^. This uncharacterized reductase can be found with either periplasmic or cytoplasmic Cld, so it is not likely to function in dissimilatory (per)chlorate reduction pathways, which are only known to occur in the periplasm ^12^. These and other co-metabolic reductases were encoded near cld in metagenomes of dryland soil, surface waters, and oxic wastewater (Figure 4C). Co-metabolic reduction of perchlorate and chlorate has long been suspected based on laboratory evidence ^8^. The genetic association of Cld with diverse enzymes with co-metabolic (per)chlorate reductase activity confirms that inadvertent reduction occurs in nature.

Most commonly, the reductases detected with Cld in metagenomes were not co-metabolic reductases but Pcr and the newly characterized group 3 Clr ^65^ (Figure 4E); other chlorate reductases were not detected. How this relates to the contribution of co-metabolic and metabolic pathways to perchlorate and chlorate reduction rates is unclear. These natural populations of dissimilatory (per)chlorate-reducing microorganisms – organisms that have had no experimental selection for the ability to respire perchlorate or chlorate – are closely related to a subset of previously identified strains and may have similar traits (Figure 4D). Curiously, the genome of one natural (per)chlorate-reducing strain, GWC2_42_11, assembled from an aquifer sediment metagenome ^66^, encodes phylogenetically divergent copies of both Pcr and group 3 Clr (Figure 4C). Two genes for Cld from this organism are found in Cld clade 6, which share a more recent ancestor with nitrite-oxidizing *Nitrospira* (clade 5, clades 7-9) than perchlorate-reducing bacteria (clade 4) (Figure 2). As a member of the class *Deltaproteobacteria* (phylum GWC2-55-46 in GTDB taxonomy), GWC2_42_11 is the most evolutionary distinct (per)chlorate-reducing bacterium identified to date, and its equally divergent reductases and Cld might help in understanding the earliest forms of perchlorate and chlorate respiration.

That ancient form of dissimilatory (per)chlorate reduction may have resembled co-metabolic (per)chlorate reduction in an organism with Cld. Cld has been shown to be inessential for removing any chlorite produced if habitats have sufficient amounts of reduced inorganic sulfur species ^67,68^ or large populations of other organisms that can degrade chlorate or chlorite ^19,61^. The above results show that chlorite stress from co-metabolic (per)chlorate reduction is a common enough phenomenon that Cld has repeatedly evolved to be co-located with co-metabolic reductase in genomes.

This is a contemporary example of how respiratory metabolisms for oxidized chlorine could have first arose from the association between chlorite dismutase and a co-metabolic reductase that later evolved to be specialized for perchlorate or chlorate reduction ^69^. If true, that might suggest that in geologic time chlorite was of consequence in biology before chlorate and perchlorate.

### Chlorite from Chlorine Oxidation

The oxidation of chlorine to chlorite is another possible reason, other than co-metabolic reduction of (per)chlorate or chemical reduction of chlorate ^70^, why chlorite affects so many diverse microorganisms in oxic habitats. A pathway for this reaction is uncertain. If it occurs, Cld should be present in organisms unable to co-metabolically reduce (per)chlorate. Using profile-HMMs representing the broad parts of the DMSO reductase family phylogeny that have perchlorate or chlorate reductase activity (PCRA) (Figure 4A), we identified genomes with Cld that do not have enzymes that reduce perchlorate or chlorate (Figure 5A). Despite the commonality of such enzymes as assimilatory nitrate reductases, this search identified 27 putative “non-(per)chlorate reducers” among isolate genomes (Supplementary Table 1). These strains represent 6 of the 19 phyla with Cld and 15 of 151 genera (Figure 5B). All are aerobes, and none were reported to be facultative anaerobes or obligate anaerobes (Figure 5B). They were isolated from diverse habitats, often characterized by high sunlight (lakes and ponds, desert rocks and sediments, growing with diatoms, cyanobacteria, or mosses) or by high amounts of reactive chlorine species (human body, wastewater treatment plant, swimming pool, showerhead biofilm) (Figure 5B). Therefore, in many habitats, the known mechanisms for the enzymatic reduction of chlorate and perchlorate appeared insufficient to explain the prevalence of chlorite and chlorite-degrading organisms.

**Figure 5.**
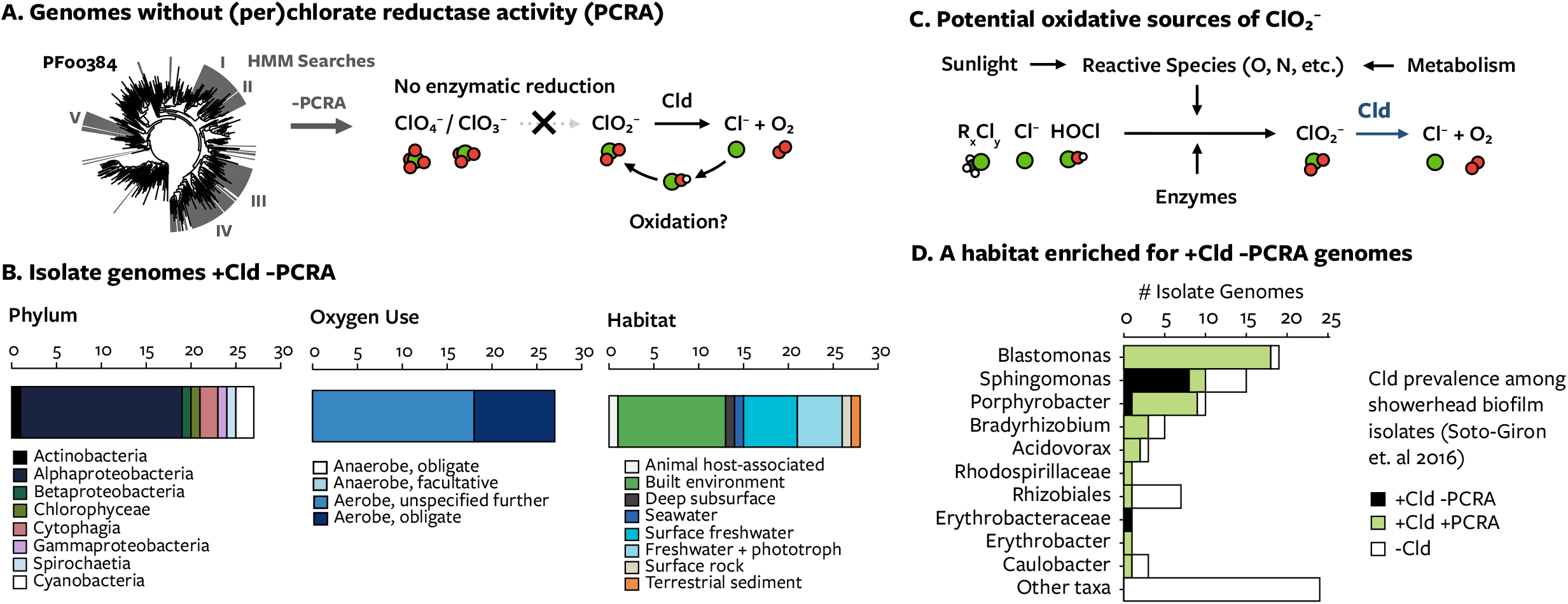
Genomes without respiratory and co-metabolic perchlorate/chlorate reductase activity (PCRA). (A) Profile-HMMs were used to find isolated microorganisms without enzymes from the broad parts of the DMSO reductase family that might have (per)chlorate reductase activity. These organisms may experience chlorite produced from oxidative chemistry. (B) The number of isolate genomes with respiratory or co-metabolic reductases grouped by phylum, relationship with oxygen, and the habitat they were isolated from (C) Pathways discussed in the text as having the potential to generate chlorite from lower oxidation states of chlorine. (D) Several organisms without PCRA were isolated from a showerhead biofilm communities exposed to chlorine residuals present in drinking water. Bars indicate the number of organisms isolated from that community with or without Cld or enzymes with putative (per)chlorate reductase activity.

One plausible route of oxidative chlorite formation is photochemistry (Figure 5C). Habitats with high sunlight are very oxidizing due to the combined effects of oxygenic phototrophy and UV photochemistry ^71,72^. Several non-(per)chlorate reducers were isolated from high sunlight habitats. UV tolerance genes were present in Cld genomic neighborhoods from several bacteria from high sunlight habitats. A putative deoxyribodipyrimidine photo-lyase, which is a light-activated protein that repairs UV-damaged DNA, is encoded in 21 Cld genomic neighborhoods and several different neighborhood groups (groups 2, 20, and 7), such as one in a betaprotebacterium in culture with *Leptolyngbya glacialis* TM1FOS73 (GCA_003242045.1) ^73^. In a sunlight photobioreactor metagenome, cld is found in four of 34 MAGs (UBA7691, UBA7681, UBA7678, UBA7677), including a *Planctomycetaceae* bacterium that has periplasmic group 2a Cld encoded near carotenoid biosynthesis genes for limiting UV photodamage ^74^. The production of chlorite from oxidative chemistry might also explain the presence of Cld in the nitrite-oxidizing bacteria (Figure 5C). Like the products of photochemistry, reactive nitrogen species nitric oxide (NO) and peroxynitrite (ONOO^-^) have high reduction potentials and can oxidize various molecules, producing other reactive species such as carbonate radicals ^75,76^.

Another plausible route of chlorine oxidation is the biochemical oxidation of hypochlorous acid to chlorite (Figure 5C). An enzyme that oxidizes hypochlorous acid to chlorite would be a major fitness benefit to organisms with Cld. Oxidation of hypochlorous acid to chlorite only requires transfer of 2 electrons and produces a less reactive product. An analogous system would be nitric oxide dioxygenase, which uses oxygen to oxidize nitric oxide to less-toxic nitrate ^77^. With Cld, oxidation of hypochlorous acid to chlorite would ultimately yield harmless chloride and oxygen. Instead of spending cellular reducing equivalents to reduce hypochlorous acid or repair oxidative damage, the enzymatic oxidation of hypochlorous acid might produce reducing equivalents. Furthermore, the removal of hypochlorous acid would limit the inhibition of Cld by hypochlorous acid ^78^. Thus, the enzymatic oxidation of hypochlorous acid to chlorite would pose major selective benefits, if it occurs.

Experimental support for this capability would be the enrichment of organisms with Cld in habitats with high hypochlorous acid. One such real-word setting appeared to be a drinking water distribution system in which of 47 of 89 strains isolated from a showerhead biofilm encoded Cld, and 10 were non-(per)chlorate reducers (Figure 5D) ^79 80^. This demonstrates a strong selection for Cld within the microbial community by the chlorine residuals present in the water. The water distribution system was expected to contain 0.8 mg/liter free residual chlorine form of hypochlorous acid and hypochlorite (ClO^-^) residuals; however, it is unclear if chlorine dioxide was used in water treatment and produced chlorite residuals (personal communication, Jorge Santo-Domingo). Except for this uncertainty, this system would meet the criteria of a habitat that selects for the ability to degrade chlorite due to only high hypochlorous acid exposure. Enzymatic oxidation of hypochlorous acid to chlorite remains an unproven hypothesis.

### A Holistic Model for Chlorine Redox Biology

The different biological processes that involve Cld suggest that the biology of chlorine reduction and oxidation should be considered as a single, bidirectional pathway (Figure 6). In this model of chlorine in biology, based both on the above genomic data and previous studies, organisms in many habitats can experience any chlorine oxyanion (HOCl, CIO_2_^-^, CIO_3_^-^, CIO_4_^-^) and some anthropogenic oxidized chlorine species (Cl_2_, ClO_2_, NH_2_Cl, NHCl_2_, and NCl_3_). Transporters appear to allow oxidized chlorine species to enter cells, where the molecules or their byproducts may be sensed and lead to changes in gene regulation.

**Figure 6.**
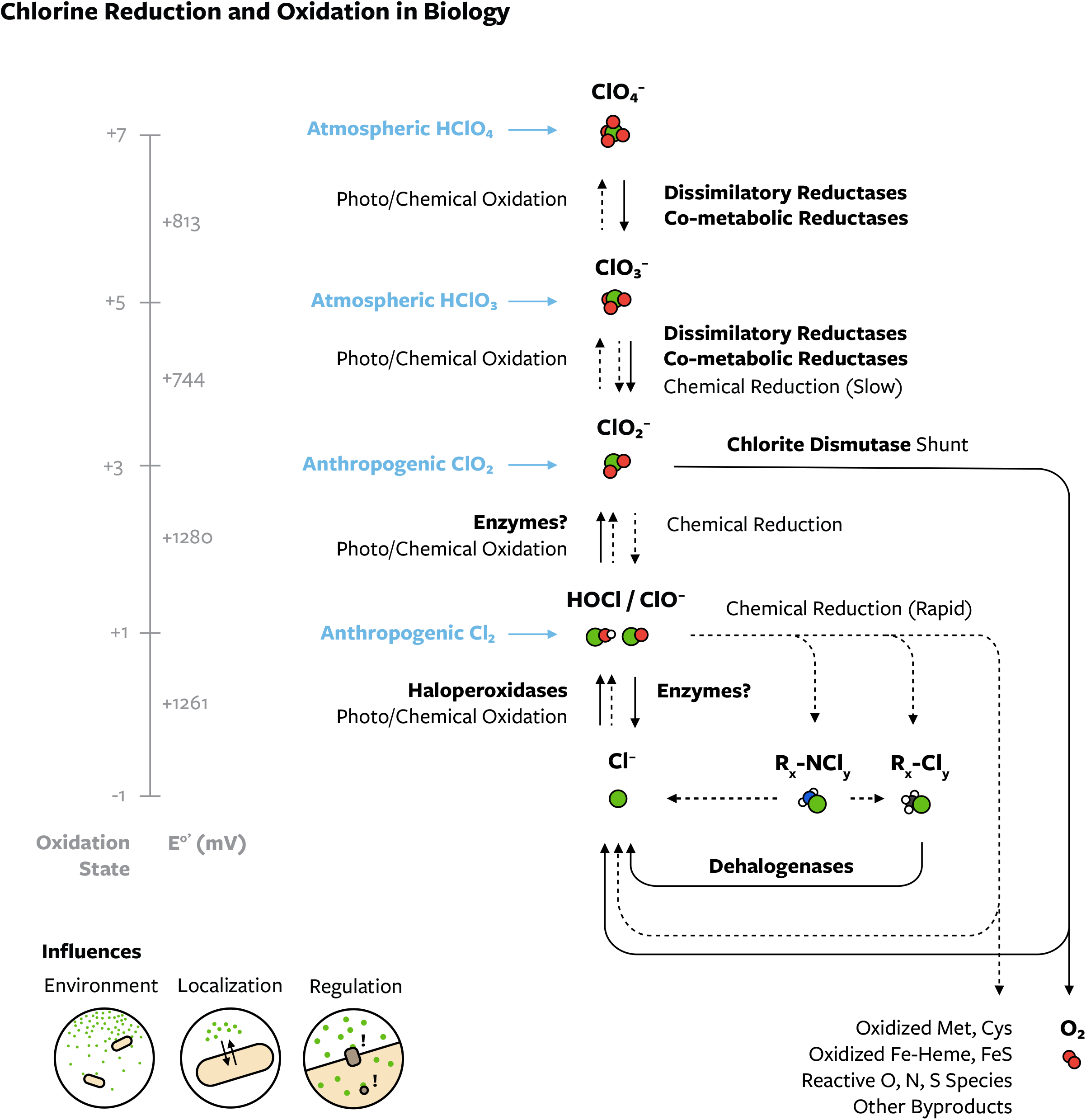
A model for biological chlorine reduction and oxidation reactions occurring within biological habitats. Biological (solid arrows, bold) and chemical or photochemical (dashed arrows) reduction and oxidation reactions of chlorine that occur, in aqueous solution or within microbial cells. Halogenases, the primary source of organochlorine, are omitted for simplicity. Other oxidized chlorine species can be external inputs into biological systems (blue). Vertical position corresponds to changes in chlorine’s formal oxidation state and reduction potential at standard conditions (pH 7, 25 °C, solutes at 1 M) in millivolts (gray). Additional factors that influence chlorine redox biology but do not perform redox reactions are shown: habitat (pH, redox potential, etc.), cellular composition including transporters, and cellular signaling and responses. Abbreviations: R-N_x_Cl_y_, organic and inorganic chloramines; R_x_-Cl_y_, organochlorine; ClO_2_, chlorine dioxide; Cl_2_, molecular chlorine.

Perchlorate, chlorate, chlorite, or hypochlorous acid, oxidized chlorine species have a propensity to be reduced in cells to the next lower oxidation state. Chlorite is produced in organisms with enzymes that can reduce perchlorate and chlorate through metabolism or co-metabolism. Cld can be considered a shunt in the reductive pathway that, when present, prevents the formation of hypochlorous acid and produces beneficial oxygen. This shunt is found in approximately 1% of microbial genomes and enriched in particular groups of bacteria.

If hypochlorous acid is formed, whether through reduction or oxidation, it reacts rapidly with biomolecules, producing a combination of chloride, chlorinated carbon and nitrogen, and oxidized byproducts ^4-6,81^. In addition to protein repair and other traditional responses to oxidative stress, a number of uncharacterized genes linked to Cld suggest a broader enzymatic response to oxidized chlorine species. A hypothetical possibility is that organisms detoxify hypochlorous acid by oxidizing hypochlorous acid to chlorite and using the chlorite dismutase shunt to degrade chlorite. In oxidizing settings, chlorite can be further oxidized (photo)chemically to chlorate or perchlorate, which are also deposited into habitats from atmosphere. The relatively stable end products of this bidirectional cycle are perchlorate and chloride. These compounds are only as inert, however, as the surrounding chemistry and biology allow.

## Conclusions

Cld, a biomarker for chlorite once thought unique to anaerobic perchlorate- and chlorate-reducing bacteria, is found in various microorganisms from both oxic and anoxic microbial habitats. This distribution suggests organisms experience significant enough amounts of chlorite in the environment to acquire Cld. The sources of chlorite are the dissimilatory reduction of (per)chlorate but also the co-metabolic reduction of (per)chlorate and, genomics suggests, the oxidation of chlorine’s lower oxidation states. That Cld participates in these pathways and in general response to reactive chlorine species justifies a model wherein oxidized chlorine species are part of a continuous, bidirectional biological pathway. Cld is subject to intermittent selection for its gain and loss, highlighting how much remains to be learned about the concentrations and fluxes of oxidized chlorine species in different environments. The expansive inventory of genes associated with Cld-encoding loci identified here provides targets for subsequent research in the biology of oxidized chlorine from regulation, transport, and repair to direct enzymatic action on chlorine-containing molecules.

## Methods

### Identification of chlorite dismutase (Cld)

A maximum likelihood phylogenetic tree of the protein family containing Cld was constructed using FastTree from the Pfam 06778 alignment of representative proteomes, at the 15% comembership threshold to limit the number of redundant proteins ^82-84^. The presence of key residues for Cld activity were identified by comparing the positions in the alignment corresponding to the distal heme arginine (R127) and proximal heme lysine (K92), histidine (H114), and glutamic acid (E167) in *Nitrobacter winogradskyii* Nb-255 ^32^. Proteins in the two major lineages of Cld were used to construct profile-hidden Markov models (HMMs) later used to identify Cld proteins ^85^. Cld and non-Cld proteins were annotated on a precomputed bacterial tree of life ^86^.

BLASTP was used to identify Cld in genomes in the JGI IMG/M, NCBI GenBank, and NCBI RefSeq databases, with RefSeq preferred ^87-89^. BLASTP was used to identify metagenomic Cld in JGI IMG/M among the largest metagenomes consisting of 90% of proteins in each “Ecosystem Category.” Metagenome-assembled genomes in the Uncultivated Bacteria and Archaea (UBA) dataset were searched directly with profile-HMMs ^90^. All Cld identified with BLASTP were confirmed with profile-HMMs. Genomic data and metadata were processed using custom scripts. Data were downloaded prior to 2020.

The fraction of a taxonomic group encoding Cld was determined by comparing the number of RefSeq genomes with the *cld* gene to the total number of RefSeq genomes available within each taxonomic group (https://github.com/kblin/ncbi-genome-download). The detection of Cld in different environments was compared using the number of cld copies per million total coding domain sequences (CDS) obtained from IMG/M metagenome metadata. Due to inconsistent definitions of environments in IMG/M, metadata were using to assign each metagenome were assigned to a custom environmental category.

### Phylogenetics

Cld proteins were aligned using MUSCLE v3.8.1551 ^91^ and built into a maximum likelihood phylogenetic tree using FastTree ^84^. The Python package ETE v. 3 was used to plot trees and to form clades of proteins at trees nodes in which the average distance to a protein was less than a selected value ^92^. N-terminal and C-terminal extensions were defined as amino acids in the alignment beyond the positions within which the average Cld protein had amino acids. Signal peptides were assigned using SignalP v. 5, accepting a positive result from any type of organism (gram-negative, gram-positive, or eukaryotic) ^93^.

Proteins in the DMSO reductase family of molybdopterin enzymes, which might function as perchlorate and chlorate reductases, were identified using a profile-HMM built from the seed alignment of Pfam 00384 ^82,85^. A maximum likelihood phylogenetic tree was constructed from those proteins encoded near *cld* and a curated set of proteins from Pfam 00384 proteins in representative proteomes at the 15% comembership threshold. Incomplete proteins were excluded using a size threshold of 300 amino acids, the size of dataset was reduced while maintaining diversity by clustering proteins at 50% amino acid identity using CD-HIT. Only positions in the alignment where a majority of proteins had residues were kept. The tree was constructed, plotted, and grouped into clades as above.

### Comparative genomics

Genes within +/- 10 positions of *cld* on the same contig were defined as part of the Cld genomic neighborhood. To compare neighborhoods, similar proteins in the neighborhood were clustered into protein subfamilies using MMSEQs v.7-4e23d set to a coverage of 0.5 and an E-value of 0.001 ^94^. The two major lineages of Cld were each defined as a separate subfamily, and subfamilies were numbered in order of their size in this dataset.

A simple statistic for gene linkage to *cld* was obtained by representing each subfamily as a node and each connection between subfamilies found in the same genomic neighborhood as edges in a network. The Python package networkx was used to compute a clustering coefficient for each node: the fraction of a node’s neighbors with an edge over the total number of edges possible between a node’s neighbors.

To simplify analysis, genomic neighborhoods with 10+ genes were grouped by similar gene content using unsupervised machine learning methods in the Python package SciKit-learn. The features of the data were the presence (1) or absence (0) of each subfamily in the neighborhood. An initial dimensional reduction was performed with Principle Components Analysis, and the resulting 50 dimensions per neighborhood were subject to t-Distributed Stochastic Neighbor Embedding (t-SNE) with a perplexity of 50 and 5,000 iterations. Neighborhoods were then clustered into groups close to each other in the two t-SNE dimensions with the Density-Based Spatial Clustering of Applications with Noise (DBSCAN) algorithm.

In select instances, genes were compared to fitness experiments using chlorite and chlorate on the Fitness Browser (fit.genomics.lbl.gov) described in Price et. al 2018.

### Data Availability

Supplementary data are available on FigShare and include: Supplementary Data 1, information on genes and genomes used in this work including accessions, taxonomy, subfamily assignments, etc. (doi:10.6084/m9.figshare.16978561); Supplementary Data 2, information on subfamilies and their clustering coefficients (doi:10.6084/m9.figshare.16980601); protein sequences found in Cld genomic neighborhoods (doi:10.6084/m9.figshare.16980613); a phylogenetic tree and alignments for Cld (doi:10.6084/m9.figshare.16982077); and profile-HMMs to identify key proteins for perchlorate, chlorate, and chlorite biology.

## Acknowledgements

This work would not be possible without the sequencing of microorganisms and microbial communities by countless scientists, for whom we are in continual gratitude. We also thank Shengqiang Shu and Joe Carlson of the Joint Genome Institute, Hirotsugu Fujitani, Jorge Santo-Domingo, and Kostas Konstantinidis, and for their personal communication regarding their work. Financial support was provided through a grant from the Energy Biosciences Institute EBI-BP program to J.D.C. and through the NSF Graduate Research Fellowship Program to T.P.B.

## Author Contributions

T.P.B. conceived of and performed all research with the guidance and supervision of J.D.C. T.P.B. and J.D.C analyzed results, wrote the manuscript, and approved its publication.

## References

1 Atashgahi, S. et al. Microbial Synthesis and Transformation of Inorganic and Organic Chlorine Compounds. Frontiers in Microbiology 9, 1–22, doi:10.3389/fmicb.2018.03079 (2018).

2 Winterton, N. Chlorine: the only green element – towards a wider acceptance of its role in natural cycles. Green Chemistry 2, 173–225, doi:10.1039/b003394o (2000).

3 Agarwal, V. et al. Enzymatic Halogenation and Dehalogenation Reactions: Pervasive and Mechanistically Diverse. Chemical Reviews, acs.chemrev.6b00571, doi:10.1021/acs.chemrev.6b00571 (2017).

4 Bengtson, P., Bastviken, D., de Boer, W. & Oberg, G. Possible role of reactive chlorine in microbial antagonism and organic matter chlorination in terrestrial environments. Environ Microbiol 11, 1330–1339, doi:10.1111/j.1462-2920.2009.01915.x (2009).

5 Winterbourn, C. C. & Kettle, A. J. Redox reactions and microbial killing in the neutrophil phagosome. Antioxid Redox Signal 18, 642–660, doi:10.1089/ars.2012.4827 (2013).

6 Comba, P., Kerscher, M., Krause, T. & Schöler, H. F. Iron-catalysed oxidation and halogenation of organic matter in nature. Environmental Chemistry 12, doi:10.1071/en14240 (2015).

7 Gray, M. J., Wholey, W.-Y. & Jakob, U. Bacterial responses to reactive chlorine species. Annual review of microbiology 67, 141–160, doi:10.1146/annurev-micro-102912-142520 (2013).

8 Youngblut, M. D., Wang, O., Barnum, T. P. & Coates, J. D. (Per)chlorate in Biology on Earth and Beyond. Annual Review of Microbiology 70, 435–459, doi:10.1146/annurev-micro-102215-095406 (2016).

9 Winterbourn, C. C. Reconciling the chemistry and biology of reactive oxygen species. Nature Chemical Biology 4, 278–286, doi:10.1038/nchembio.85 (2008).

10 Deborde, M. & von Gunten, U. Reactions of chlorine with inorganic and organic compounds during water treatment - Kinetics and mechanisms: A critical review. Water Research 42, 13–51, doi:10.1016/j.watres.2007.07.025 (2008).

11 Asami, M., Kosaka, K. & Kunikane, S. Bromate, chlorate, chlorite and perchlorate in sodium hypochlorite solution used in water supply. Journal of Water Supply: Research and Technology - AQUA 58, 107–115, doi:10.2166/aqua.2009.014 (2009).

12 Coates, J. D. & Achenbach, L. A. Microbial perchlorate reduction: rocket-fueled metabolism. Nature Reviews Microbiology 2, 569–580, doi:10.1038/nrmicro926 (2004).

13 Leri, A. C. et al. A marine sink for chlorine in natural organic matter. Nature Geoscience 8, 620–624, doi:10.1038/NGEO2481 (2015).

14 Ortiz-Bermudez, P., Hirth, K. C., Srebotnik, E. & Hammel, K. E. Chlorination of lignin by ubiquitous fungi has a likely role in global organochlorine production. PNAS 104, 3895–3900 (2007).

15 Rao, B. et al. Perchlorate production by photodecomposition of aqueous chlorine solutions. Environ Sci Technol 46, 11635–11643, doi:10.1021/es3015277 (2012).

16 Rajagopalan, S. et al. Perchlorate in wet deposition across North America. Environmental Science and Technology 43, 616–622, doi:10.1021/es801737u (2009).

17 Dasgupta, P. K. et al. The origin of naturally occurring perchlorate: The role of atmospheric processes. Environmental Science and Technology 39, 1569–1575, doi:10.1021/es048612x (2005).

18 Youngblut, M. D. et al. Perchlorate reductase is distinguished by active site aromatic gate residues. Journal of Biological Chemistry 291, 9190–9202, doi:10.1074/jbc.M116.714618 (2016).

19 Barnum, T. P. et al. Identification of a parasitic symbiosis between respiratory metabolisms in the biogeochemical chlorine cycle. The ISME Journal, doi:10.1038/s41396-020-0599-1 (2020).

20 Leimkühler, S. & Iobbi-Nivol, C. Bacterial molybdoenzymes: old enzymes for new purposes. FEMS Microbiology Reviews 40, 1–18, doi:10.1093/femsre/fuv043 (2015).

21 Weiner, J. H., MacIsaac, D. P., Bishop, R. E. & Bilous, P. T. Purification and properties of Escherichia coli dimethyl sulfoxide reductase, an iron-sulfur molybdoenzyme with broad substrate specificity. 170, 1505–1510, doi:10.1128/jb.170.4.1505-1510.1988 %J Journal of Bacteriology (1988).

22 Hinsley, A. P. & Berks, B. C. Specificity of respiratory pathways involved in the reduction of sulfur compounds by Salmonella enterica. 148, 3631–3638, doi:https://doi.org/10.1099/00221287-148-11-3631 (2002).

23 Riggs, D. L., Tang, J. S. & Barrett, E. L. Thiosulfate reductase as a chlorate reductase in Salmonella typhimurium. FEMS Microbiology Letters 44, 427–430, doi:10.1111/j.1574-6968.1987.tb02326.x %J FEMS Microbiology Letters (1987).

24 McEwan, A. G., Wetzstein, H. G., Meyer, O., Jackson, J. B. & Ferguson, S. J. The periplasmic nitrate reductase of Rhodobacter capsulatus; purification, characterisation and distinction from a single reductase for trimethylamine-N-oxide, dimethylsulphoxide and chlorate. Archives of Microbiology 147, 340–345 (1987).

25 Celis, A. I. et al. A dimeric chlorite dismutase exhibits O2-generating activity and acts as a chlorite antioxidant in Klebsiella pneumoniae MGH 78578. Biochemistry 54, 434–446, doi:10.1021/bi501184c (2015).

26 Schaffner, I. et al. Mechanism of chlorite degradation to chloride and dioxygen by the enzyme chlorite dismutase. Archives of Biochemistry and Biophysics 574, 18–26, doi:10.1016/j.abb.2015.02.031 (2015).

27 Van Ginkel, C. G., Rikken, G. B., Kroon, A. G. M. & Kengen, S. W. M. Purification and characterization of chlorite dismutase: A novel oxygen-generating enzyme. Archives of Microbiology 166, 321–326, doi:10.1007/s002030050390 (1996).

28 Dubois, J. L. O – O Bond Formation by a Heme Protein : The Unexpected Efficiency of Chlorite Dismutase. Molecular Water Oxidation Catalysis: A Key Topic for New Sustainable Energy Conversion Schemes (2014).

29 Rikken, G. B., Kroon, A. G. M. & Van Ginkel, C. G. Transformation of (per)chlorate into chloride by a newly isolated bacterium: reduction and dismutation. Applied Microbiology and Biotechnology 45, 420–426, doi:10.1007/s002530050707 (1996).

30 Coates, J. D. et al. Ubiquity and diversity of dissimilatory (per)chlorate-reducing bacteria. Applied and Environmental Microbiology 65, 5234–5241 (1999).

31 Maixner, F. et al. Environmental genomics reveals a functional chlorite dismutase in the nitrite-oxidizing bacterium ‘Candidatus Nitrospira defluvii’. Environmental Microbiology 10, 3043–3056, doi:10.1111/j.1462-2920.2008.01646.x (2008).

32 Hofbauer, S. et al. From chlorite dismutase towards HemQ-the role of the proximal H-bonding network in haeme binding. Bioscience reports 36, e00312, doi:10.1042/BSR20150330 (2016).

33 de Geus, D. C. et al. Crystal Structure of Chlorite Dismutase, a Detoxifying Enzyme Producing Molecular Oxygen. Journal of Molecular Biology 387, 192–206, doi:https://doi.org/10.1016/j.jmb.2009.01.036 (2009).

34 Celis, A. I. & DuBois, J. L. Substrate, product, and cofactor: The extraordinarily flexible relationship between the CDE superfamily and heme. Archives of Biochemistry and Biophysics 574, 3–17, doi:10.1016/j.abb.2015.03.004 (2015).

35 Zámocký, M. et al. Independent evolution of four heme peroxidase superfamilies. Archives of Biochemistry and Biophysics 574, 108–119, doi:10.1016/j.abb.2014.12.025 (2015).

36 Dailey, H.a. & Gerdes, S. HemQ: An iron-coproporphyrin oxidative decarboxylase for protoheme synthesis in Firmicutes and Actinobacteria. Archives of Biochemistry and Biophysics 574, 27–35, doi:10.1016/j.abb.2015.02.017 (2015).

37 Schaffner, I. et al. Dimeric chlorite dismutase from the nitrogen-fixing cyanobacterium Cyanothece sp. PCC7425. Molecular Microbiology 96, 1053–1068, doi:10.1111/mmi.12989 (2015).

38 Mlynek, G. et al. Unexpected Diversity of Chlorite Dismutases: a Catalytically Efficient Dimeric Enzyme from Nitrobacter winogradskyi. 193, 2408–2417, doi:10.1128/JB.01262-10 (2011).

39 Bogen, C. et al. Reconstruction of the lipid metabolism for the microalga Monoraphidium neglectum from its genome sequence reveals characteristics suitable for biofuel production. BMC Genomics 14, 926, doi:10.1186/1471-2164-14-926 (2013).

40 Cary, S. C., McDonald, I. R., Barrett, J. E. & Cowan, D. A. On the rocks: the microbiology of Antarctic Dry Valley soils. Nat Rev Microbiol 8, 129–138, doi:10.1038/nrmicro2281 (2010).

41 Maccario, L., Carpenter, S. D., Deming, J. W., Vogel, T. M. & Larose, C. Sources and selection of snow-specific microbial communities in a Greenlandic sea ice snow cover. Sci Rep 9, 2290, doi:10.1038/s41598-019-38744-y (2019).

42 Dubois, J. L. & Ojha, S. Sustaining Life on Planet Earth: Metalloenzymes Mastering Dioxygen and Other Chewy Gases. 15, 45–87, doi:10.1007/978-3-319-12415-5 (2015).

43 Lucker, S., Nowka, B., Rattei, T., Spieck, E. & Daims, H. The Genome of Nitrospina gracilis Illuminates the Metabolism and Evolution of the Major Marine Nitrite Oxidizer. Front Microbiol 4, 27, doi:10.3389/fmicb.2013.00027 (2013).

44 Enright, A. J., Iliopoulos, I., Kyrpides, N. C. & Ouzounis, C. A. Protein interaction maps for complete genomes based on gene fusion events. Nature 402, 86–90, doi:10.1038/47056 (1999).

45 Kitzinger, K. et al. Characterization of the First “Candidatus Nitrotoga” Isolate Reveals Metabolic Versatility and Separate Evolution of Widespread Nitrite-Oxidizing Bacteria. MBio 9, e01186–01118, doi:10.1128/mBio.01186-18 %J mBio (2018).

46 Thandar, S. M., Ushiki, N., Fujitani, H., Sekiguchi, Y. & Tsuneda, S. Ecophysiology and Comparative Genomics of Nitrosomonas mobilis Ms1 Isolated from Autotrophic Nitrifying Granules of Wastewater Treatment Bioreactor. Frontiers in Microbiology 7, 1869, doi:10.3389/fmicb.2016.01869 (2016).

47 Fujitani, H., Kumagai, A., Ushiki, N., Momiuchi, K. & Tsuneda, S. Selective isolation of ammonia-oxidizing bacteria from autotrophic nitrifying granules by applying cell-sorting and sub-culturing of microcolonies. Frontiers in Microbiology 6, 1–10, doi:10.3389/fmicb.2015.01159 (2015).

48 Overbeek, R., Fonstein, M., D’Souza, M., Gordon, D. P. & Maltsev, N. The use of gene clusters to infer functional coupling. Proc Natl Acad Sci USA 96, 2896–2901 (1999).

49 Pellegrini, M., Marcotte, E., Thompson, M. J., Eisenberg, D. & Yeates, T. O. Assigning protein functions by comparative genome analysis: Protein phylogenetic profiles. Proc Natl Acad Sci USA 96, 4285–4288 (1999).

50 Melnyk, R. a. et al. Identification of a perchlorate reduction genomic island with novel regulatory and metabolic genes. Applied and environmental microbiology 77, 7401–7404, doi:10.1128/AEM.05758-11 (2011).

51 Clark, I. C., Melnyk, R. a., Engelbrektson, A. & Coates, J. D. Structure and evolution of chlorate reduction composite transposons. mBio 4, e00379–00313, doi:10.1128/mBio.00379-13 (2013).

52 Melnyk, R. A. & Coates, J. D. The Perchlorate Reduction Genomic Island: Mechanisms and Pathways of Evolution by Horizontal Gene Transfer. BMC Genomics 16, 862, doi:10.1186/s12864-015-2011-5 (2015).

53 Nontaleerak, B. et al. Roles of RcsA, an AhpD Family Protein, in Reactive Chlorine Stress Resistance and Virulence in Pseudomonas aeruginosa. Applied and Environmental Microbiology 86, e01480–01420, doi:10.1128/AEM.01480-20 (2020).

54 Kallberg, Y., Oppermann, U. & Persson, B. Classification of the short-chain dehydrogenase/reductase superfamily using hidden Markov models. FEBS J 277, 2375–2386, doi:10.1111/j.1742-4658.2010.07656.x (2010).

55 Price, M. N. et al. Mutant phenotypes for thousands of bacterial genes of unknown function. Nature 557, 503–509, doi:10.1038/s41586-018-0124-0 (2018).

56 Melnyk, R. a. et al. Novel Mechanism for Scavenging of Hypochlorite Involving a Periplasmic Methionine-Rich Peptide and Methionine Sulfoxide Reductase. mBio 6, e00233–00215, doi:10.1128/mBio.00233-15.Editor (2015).

57 Sultana, S., Foti, A. & Dahl, J.-U. Bacterial Defense Systems against the Neutrophilic Oxidant Hypochlorous Acid. Infection and Immunity 88, e00964–00919, doi:10.1128/IAI.00964-19 (2020).

58 Stewart, V. Nitrate respiration in relation to facultative metabolism in enterobacteria. Microbiol Rev 52, 190–232 (1988).

59 Tran, P. et al. Microbial life under ice: Metagenome diversity and in situ activity of Verrucomicrobia in seasonally ice-covered Lakes. Environmental Microbiology 20, 2568–2584, doi:https://doi.org/10.1111/1462-2920.14283 (2018).

60 Tsuji, J. M. et al. Anoxygenic photosynthesis and iron–sulfur metabolic potential of Chlorobia populations from seasonally anoxic Boreal Shield lakes. The ISME Journal 14, 2732–2747, doi:10.1038/s41396-020-0725-0 (2020).

61 Clark, I. C. et al. Genetic dissection of chlorate respiration in Pseudomonas stutzeri PDA reveals syntrophic (per)chlorate reduction. Environmental Microbiology 18, 3342–3354, doi:10.1111/1462-2920.13068 (2016).

62 Chapman, A. L. P., Hampton, M. B., Senthilmohan, R., Winterbourn, C. C. & Kettle, A. J. Chlorination of Bacterial and Neutrophil Proteins during Phagocytosis and Killing of Staphylococcus aureus. Journal of Biological Chemistry 277, 9757–9762, doi:https://doi.org/10.1074/jbc.M106134200 (2002).

63 Liu, Q., Song, W.-Z., Zhou, Y.-G., Dong, X.-Z. & Xin, Y.-H. Phenotypic divergence of thermotolerance: Molecular basis and cold adaptive evolution related to intrinsic DNA flexibility of glacier-inhabiting Cryobacterium strains. Environmental Microbiology 22, 1409–1420, doi:https://doi.org/10.1111/1462-2920.14957 (2020).

64 Singh, D., Kumari, A., Ramaswamy, S. & Ramanathan, G. Expression, purification and substrate specificities of 3-nitrotoluene dioxygenase from Diaphorobacter sp. strain DS2. Biochemical and Biophysical Research Communications 445, 36–42, doi:https://doi.org/10.1016/j.bbrc.2014.01.113 (2014).

65 Barnum, T. P. & Coates, J. D. An uncharacterized clade in the DMSO reductase family of molybdenum oxidoreductases is a new type of chlorate reductase. Environ Microbiol Rep 12, 534–539, doi:10.1111/1758-2229.12869 (2020).

66 Anantharaman, K. et al. Thousands of microbial genomes shed light on interconnected biogeochemical processes in an aquifer system. Nature communications 7, 13219, doi:10.1038/ncomms13219 (2016).

67 Liebensteiner, M. G. et al. Perchlorate and chlorate reduction by the Crenarchaeon Aeropyrum pernix and two thermophilic Firmicutes. Environmental Microbiology Reports 7, 936–945, doi:10.1111/1758-2229.12335 (2015).

68 Liebensteiner, M. G., Pinkse, M. W. H., Schaap, P. J., Stams, A. J. M. & Lomans, B. P. Archaeal (per)chlorate reduction at high temperature: an interplay of biotic and abiotic reactions. Science (New York, N.Y.) 340, 85–87, doi:10.1126/science.1233957 (2013).

69 Barnum, T. P. et al. Genome-resolved metagenomics identifies genetic mobility, metabolic interactions, and unexpected diversity in perchlorate-reducing communities. ISME Journal 12, 1568–1581, doi:10.1038/s41396-018-0081-5 (2018).

70 Brundrett, M., Yan, W., Velazquez, M. C., Rao, B. & Jackson, W. A. Abiotic Reduction of Chlorate by Fe(II) Minerals: Implications for Occurrence and Transformation of Oxy-Chlorine Species on Earth and Mars. ACS Earth and Space Chemistry 3, 700–710, doi:10.1021/acsearthspacechem.8b00206 (2019).

71 Gao, Q. & Garcia-Pichel, F. Microbial ultraviolet sunscreens. Nat Rev Microbiol 9, 791–802, doi:10.1038/nrmicro2649 (2011).

72 Latifi, A., Ruiz, M. & Zhang, C. C. Oxidative stress in cyanobacteria. FEMS Microbiol Rev 33, 258–278, doi:10.1111/j.1574-6976.2008.00134.x (2009).

73 Cornet, L. et al. Metagenomic assembly of new (sub)polar Cyanobacteria and their associated microbiome from non-axenic cultures. Microb Genom 4, doi:10.1099/mgen.0.000212 (2018).

74 Krohn-Molt, I. et al. Metagenome survey of a multispecies and alga-associated biofilm revealed key elements of bacterial-algal interactions in photobioreactors. Appl Environ Microbiol 79, 6196–6206, doi:10.1128/AEM.01641-13 (2013).

75 Radi, R. Oxygen radicals, nitric oxide, and peroxynitrite: Redox pathways in molecular medicine. Proc Natl Acad Sci. USA 115, 5839–5848, doi:10.1073/pnas.1804932115 (2018).

76 Moller, M. N. et al. Detection and quantification of nitric oxide-derived oxidants in biological systems. J Biol Chem 294, 14776–14802, doi:10.1074/jbc.REV119.006136 (2019).

77 Forrester, M. T. & Foster, M. W. Protection from nitrosative stress: a central role for microbial flavohemoglobin. Free Radic Biol Med 52, 1620–1633, doi:10.1016/j.freeradbiomed.2012.01.028 (2012).

78 Hofbauer, S. et al. Transiently produced hypochlorite is responsible for the irreversible inhibition of chlorite dismutase. Biochemistry 53, 3145–3157, doi:10.1021/bi500401k (2014).

79 Soto-Giron, M. J. et al. Biofilms on Hospital Shower Hoses: Characterization and Implications for Nosocomial Infections. Appl Environ Microbiol 82, 2872–2883, doi:10.1128/AEM.03529-15 (2016).

80 Le Dantec, C. et al. Chlorine disinfection of atypical mycobacteria isolated from a water distribution system. Appl Environ Microbiol 68, 1025–1032, doi:10.1128/aem.68.3.1025-1032.2002 (2002).

81 Hwang, C., Ling, F., Andersen, G. L., LeChevallier, M. W. & Liu, W. T. Microbial community dynamics of an urban drinking water distribution system subjected to phases of chloramination and chlorination treatments. Appl Environ Microbiol 78, 7856–7865, doi:10.1128/AEM.01892-12 (2012).

82 El-Gebali, S. et al. The Pfam protein families database in 2019. Nucleic Acids Research 47, D427–D432, doi:10.1093/nar/gky995 %J Nucleic Acids Research (2018).

83 Chen, C. et al. Representative Proteomes: A Stable, Scalable and Unbiased Proteome Set for Sequence Analysis and Functional Annotation. PLOS ONE 6, e18910, doi:10.1371/journal.pone.0018910 (2011).

84 Price, M. N., Dehal, P. S. & Arkin, A. P. J. P. o. FastTree 2–approximately maximum-likelihood trees for large alignments. 5, e9490 (2010).

85 Finn, R. D. et al. HMMER web server: 2015 update. Nucleic Acids Research, 1–9, doi:10.1093/nar/gkv397 (2015).

86 Mendler, K. et al. AnnoTree: visualization and exploration of a functionally annotated microbial tree of life. Nucleic Acids Research 47, 4442–4448, doi:10.1093/nar/gkz246 %J Nucleic Acids Research (2019).

87 Camacho, C. et al. BLAST plus: architecture and applications. BMC Bioinformatics 10, 421, doi:Artn 421\nDoi 10.1186/1471-2105-10-421 (2009).

88 Chen, I.-M. A. et al. IMG/M: integrated genome and metagenome comparative data analysis system. Nucleic acids research 45, 507–516, doi:10.1093/nar/gkw929 (2017).

89 Coordinators, N. R., Om-, G. E., Viewer, M. & Read, S. Database resources of the National Center for Biotechnology Information. Nucleic acids research 43, 6–17, doi:10.1093/nar/gku1130 (2014).

90 Parks, D. H. et al. Recovery of nearly 8,000 metagenome-assembled genomes substantially expands the tree of life. Nature Microbiology 2, 1533–1542, doi:10.1038/s41564-017-0012-7 (2017).

91 Edgar, R. C. MUSCLE: multiple sequence alignment with high accuracy and high throughput. Nucleic acids research 32, 1792–1797, doi:10.1093/nar/gkh340 (2004).

92 Huerta-Cepas, J., Serra, F. & Bork, P. ETE 3: Reconstruction, Analysis, and Visualization of Phylogenomic Data. Molecular Biology and Evolution 33, 1635–1638, doi:10.1093/molbev/msw046 %J Molecular Biology and Evolution (2016).

93 Almagro Armenteros, J. J. et al. SignalP 5.0 improves signal peptide predictions using deep neural networks. Nature Biotechnology 37, 420–423, doi:10.1038/s41587-019-0036-z (2019).

94 Hauser, M., Steinegger, M. & Söding, J. J. B. MMseqs software suite for fast and deep clustering and searching of large protein sequence sets. 32, 1323–1330 (2016).

