## Supplementary information for "Chlorine redox chemistry is not rare in biology"

**Supplementary Figure 1**
Maximum likelihood phylogeny of Cld lineage 2.

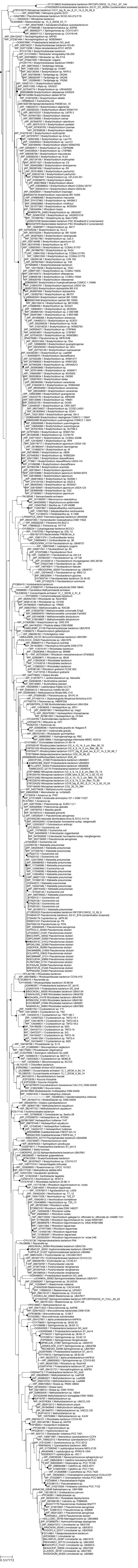

**Supplementary Figure 2**

Maximum likelihood tree of Cld lineage 1.

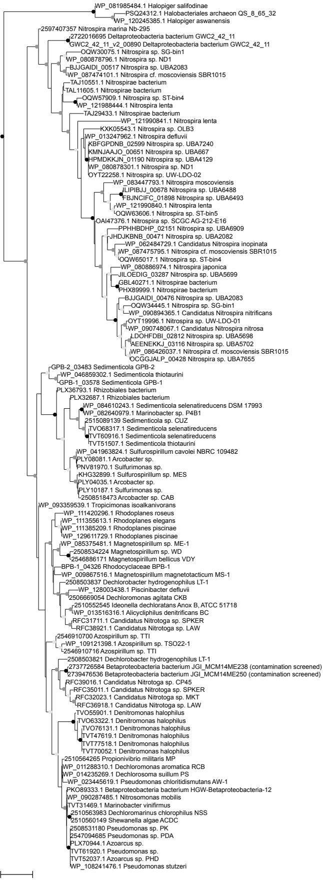

### Supplementary Figure 3

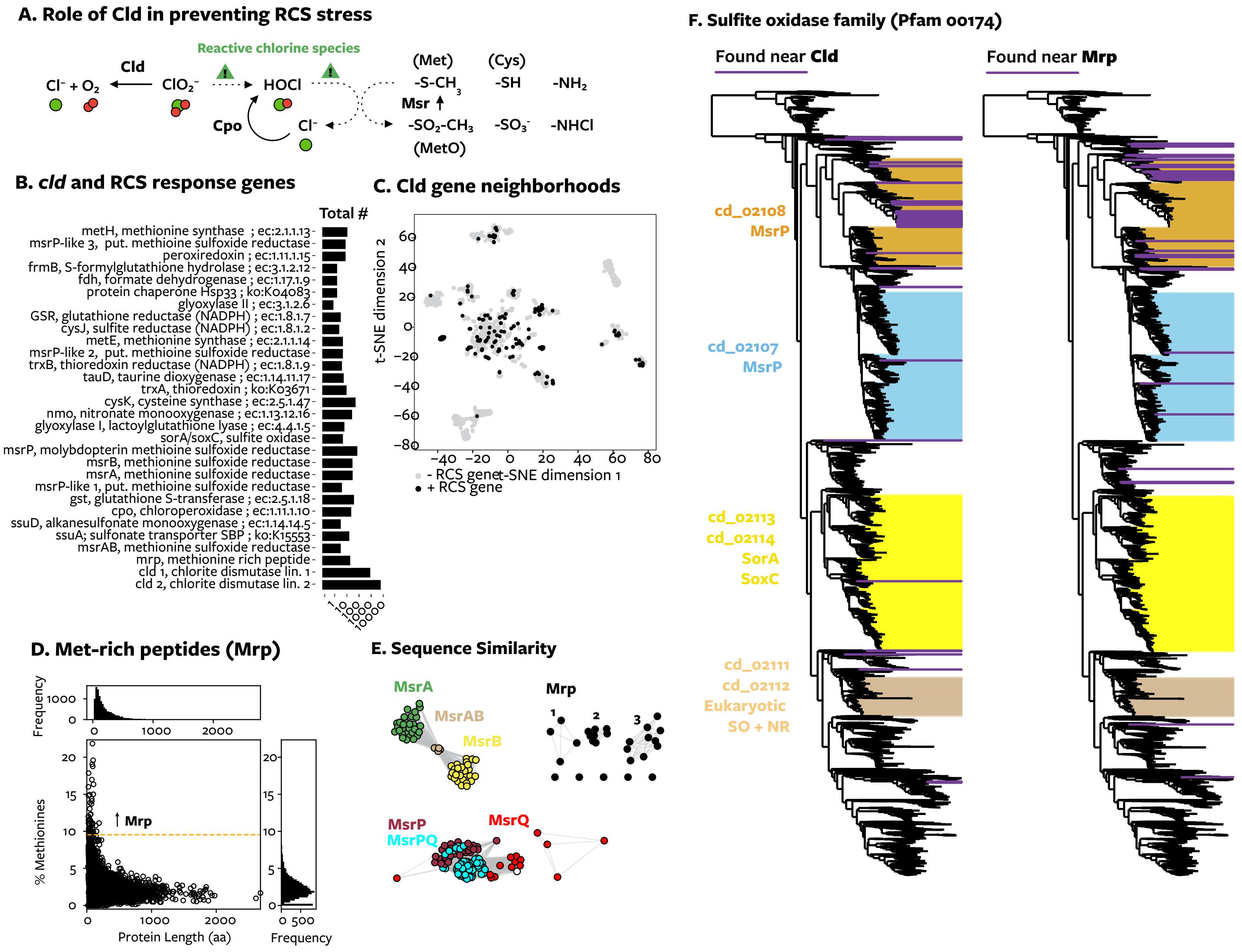

**Description of Supplementary Figure 3**

This figure shows the association of Cld with genes for reactive chlorine species response.

Panel A is a schematic diagram illustrating why Cld, having chlorite as a substrate, would be associated with HOCl and genes for repairing HOCl-mediated oxidative damage: HOCl is produced from the reduction of chlorite and may be a major source of oxidative damage in chlorite reduction. Panel B shows the frequency of reactive chlorine species response genes found with Cld, in number of genes detected. Panel C shows the Cld genomic neighborhoods including reactive chlorine species response genes. Together these data show how widespread known genes for reactive chlorine species response are within Cld genomic neighborhoods.

An important component of reactive chlorine species response in need of further definition is the methionine sulfoxide reductase system. Hypochlorous acid most rapidly and specifically oxidizes sulfur atoms in amino acids, converting methionine to methionine sulfoxide and progressively oxidizing cysteine to sulfenic acid [1,2]. Methionine is regenerated by methionine sulfoxide reductases: MsrA, MsrB, and MsrP (formerly YedY) [3-5]. The importance of MsrP reductases is evident from their being the most common beneficial genes in chlorite stress conditions [6], but MsrP and Mrp – a methionine rich peptides that scavenges hypochlorous acid and chlorite is sometimes co-expressed with MsrP [5] – are poorly defined.

New putative Mrp were defined by fitting a normal distribution to the mean Met content of all protein subfamilies in Cld genomic neighborhoods and selecting small proteins with exceptional Met content ($p< 0.00001$, >8.7% Met) found with MsrA, MsrB, or MsrP. Panel D shows the distribution of the mean methionine content in protein subfamilies (% methionine) used to define methionine rich peptides (Mrp). The phylogenetic relationships between proteins in the methionine sulfoxide reductase system were investigated using a network analysis, shown in panel E. Unlike Msr enzymes, Mrp formed multiple distinct groups that had no significant sequence similarity to one another, indicating these short HOCl-scavenging peptides might have evolved independently multiple times. These peptides may be like other short peptides that evolved *de novo*, from noncoding sequences [7].

MsrP methionine sulfoxide reductases belong to a larger family of proteins, the sulfite oxidase family of molybdopterin enzymes (Pfam 00174), which has several conserved domains of unknown function. To define which proteins in the sulfite oxidase family act as methionine sulfoxide reductases, proteins found with Cld were placed into a maximum likelihood phylogenetic tree of Pfam 00174 including proteins from representative proteomes. Extending a previous analysis using Mrp [5], nodes in the tree were further annotated if Mrp, a substrate of MsrP, was found within 5 genes of the molybdopterin domain gene. Panel F shows a version of the tree with each annotation: proteins found with Cld (left tree) or Mrp (right tree) are highlighted blue, and clades are highly by their conserved domain annotation. These genomic signatures of activity span the breadth of both conserved domains that contain characterized methionine sulfoxide reductases: MsrP from *Azospira suillum* PS (cd_02108) and MsrP from *Escherichia coli* and *Rhodobacter sphaeroides* (cd_02107) [4,5,8]. The occurrence of Cld and Mrp were found in other clades that could be methionine sulfoxide reductase or are traditional sulfite oxidases. This could be spurious, but a potential functional link between sulfite oxidases and Cld may be related to the involvement of sulfite oxidases in recycling sulfur oxidized by reactive oxidants [9].


1 Gray, M. J., Wholey, W.-Y. & Jakob, U. Bacterial responses to reactive chlorine species. *Annual review of microbiology* **67**, 141-160, doi:10.1146/annurev-micro-102912-142520 (2013).

2 Winterbourn, C. C. Reconciling the chemistry and biology of reactive oxygen species. *Nature Chemical Biology* **4**, 278-286, doi:10.1038/nchembio.85 (2008).

3 Ezraty, B., Gennaris, A., Barras, F. & Collet, J.-F. Oxidative stress, protein damage and repair in bacteria. *Nature Reviews Microbiology* **15**, 385 (2017).

4 Gennaris, A. *et al.* Repairing oxidized proteins in the bacterial envelope using respiratory chain electrons. *Nature* **528**, 409-412, doi:10.1038/nature15764 (2015).

5 Melnyk, R. a. *et al.* Novel Mechanism for Scavenging of Hypochlorite Involving a Periplasmic Methionine-Rich Peptide and Methionine Sulfoxide Reductase. *mBio* **6**, e00233-00215, doi:10.1128/mBio.00233-15.Editor (2015).

6 Price, M. N. *et al.* Mutant phenotypes for thousands of bacterial genes of unknown function. *Nature* **557**, 503-509, doi:10.1038/s41586-018-0124-0 (2018).

7 Knopp, M. *et al.* De Novo Emergence of Peptides That Confer Antibiotic Resistance. **10**, e00837-00819, doi:10.1128/mBio (2019).

8 Tarrago, L. *et al.* Rhodobacter sphaeroides methionine sulfoxide reductase P reduces R- and S-diastereomers of methionine sulfoxide from a broad-spectrum of protein substrates. *Biochem J* **475**, 3779-3795, doi:10.1042/BCJ20180706 (2018).

9 Kappler, U. & Schwarz, G. The Sulfite Oxidase Family of Molybdenum Enzymes. doi:10.1039/9781782620891-00240 (2017).
